# Pharmacological inhibition of V-ATPase targets mode-switching but not the proton transport cycle

**DOI:** 10.1101/2025.09.28.679038

**Authors:** Eleftherios Kosmidis, Maria Fokaeos, Christopher G. Shuttle, Michael C. Isselstein, Aleksander Cvjetkovic, Peter J. Johnson, Jesper L. Pederesen, Reinhard Jahn, Julia Preobraschenski, Dimitrios Stamou

## Abstract

Vacuolar-type adenosine triphosphatases (V-ATPases) are rotary proton pumps that establish proton gradients across cellular membranes^1,2^. Their pharmacological inhibition is currently under active investigation as a therapeutic strategy for cancer, infectious diseases, and autophagy-related disorders^3,4^. However, the molecular mechanism underlying V-ATPase inhibition remains poorly understood. Based on ensemble average measurements, it is widely assumed that inhibitors suppress activity by slowing the catalytic transport cycle and reducing proton transport rates^5–7^. Here, we tested this popular notion by directly measuring single-molecule proton pumping in the presence of three potent V-ATPase inhibitors: bafilomycin A1, concanamycin A, and diphyllin. Although all compounds abolish proton gradients in a canonical concentration-dependent manner (IC_50_ of 0.2 nM, 0.6 nM, and 41 nM, respectively), they leave the proton transport rate of active V-ATPases essentially unchanged. Instead, inhibitors modulate the reversible switching kinetics between ultralong-lived active (pumping) and inactive modes. Distinct inhibitors modulate mode lifetimes in a mode-specific and differentially efficient manner, altering the probability of the pump being in the active mode. Given that mode-switching has been documented across diverse primary^8,9^ and secondary^10–13^ active transporters, our results suggest a novel strategy for therapeutic intervention that targets mode occupancy rather than the canonical transport cycle.

## Introduction

The vacuolar-type ATPase (V-ATPase) is a multi-subunit rotary mechanoenzyme essential for acidifying intracellular compartments in all eukaryotic cells^2,14,15^. Structurally, the V-ATPase comprises two main domains^16^. The soluble V_1_ domain, located in the cytoplasm, is responsible for ATP hydrolysis, while the membrane-integral V_O_ domain shuttles protons across the bilayer^16^. The coupling of the two domains facilitates the ATP hydrolysis-driven pumping of protons (H⁺ ions) across various membranes, resulting in the acidification of organelles such as lysosomes, endosomes, the Golgi apparatus, synaptic and secretory vesicles^2^.

Acid-base homeostasis is critical for numerous cellular processes; therefore, overactivity of the V-ATPase is increasingly recognized as a driver of various pathophysiological conditions^4^. For example, in cancer, it acidifies the tumour microenvironment, promoting invasion, metastasis, and chemotherapy resistance^3^. In osteoporosis, excessive V-ATPase activity accelerates bone resorption^17^, while pathogenic microorganisms exploit V-ATPase to enhance survival. Moreover, V-ATPase activity has direct implications for the cytosolic delivery of macromolecular therapeutics which often relies on mechanisms sensitive to endosomal pH^18,19^. Given these diverse roles, the V-ATPase represents a promising pharmacological target^20^.

Pharmacological inhibition of V-ATPases is widely assumed to reduce the proton transport rates by targeting the proton transport cycle^5–7^. Here, we directly tested this hypothesis through single-molecule measurements of proton pumping^9^. Our investigation was motivated by recent findings that the mammalian V-ATPase is not continuously active over time, instead, it can stochastically switch between ultralong-lived, physiologically relevant, active and inactive modes^8^. Importantly, regulation of proton pumping can occur either through the proton transport cycle or by tuning the switching probability between the active and inactive modes^8^. Here, direct observation of proton pumping allowed us to deconvolute these two processes and delineate the molecular mechanism of pharmacological inhibition. Intriguingly, experiments in the presence of three potent inhibitors (IC_50_ in the nM range) demonstrate that V-ATPase inhibition does not require modulation of the canonical proton transport cycle but can instead be exclusively mediated through mode switching.

### Mode-switching by single V-ATPases in single native synaptic vesicles (SVs)

We monitored proton pumping by single V-ATPases in single intact synaptic vesicles (SVs)^8^ isolated and purified from rat brain^21,22^ (Fig. 1, Supp. Fig. 1), leveraging a recently established and validated methodology^8,9^. In Figure 1, we provide a concise outline of the method^8,9^, along with the key observables and findings. Briefly, intact SVs were labelled with a photostable pH-sensitive lipid-conjugated fluorophore (pHrodo-DOPE)^8,9^, tethered on a passivated surface at low densities, and imaged over time at the single vesicle-level^23–26^ using fluorescence microscopy (Fig. 1a, b). Because SVs predominantly carry only one copy of the V-ATPase^8,27–29^ the ATP-dependent changes in the intensity of single SVs are directly proportional to the pumping of protons by single V-ATPases into the SV lumen^8^ (Fig. 1d, E.D. Fig. 1, Supp. Fig. 2, Supplementary Methods).

**Fig. 1.**
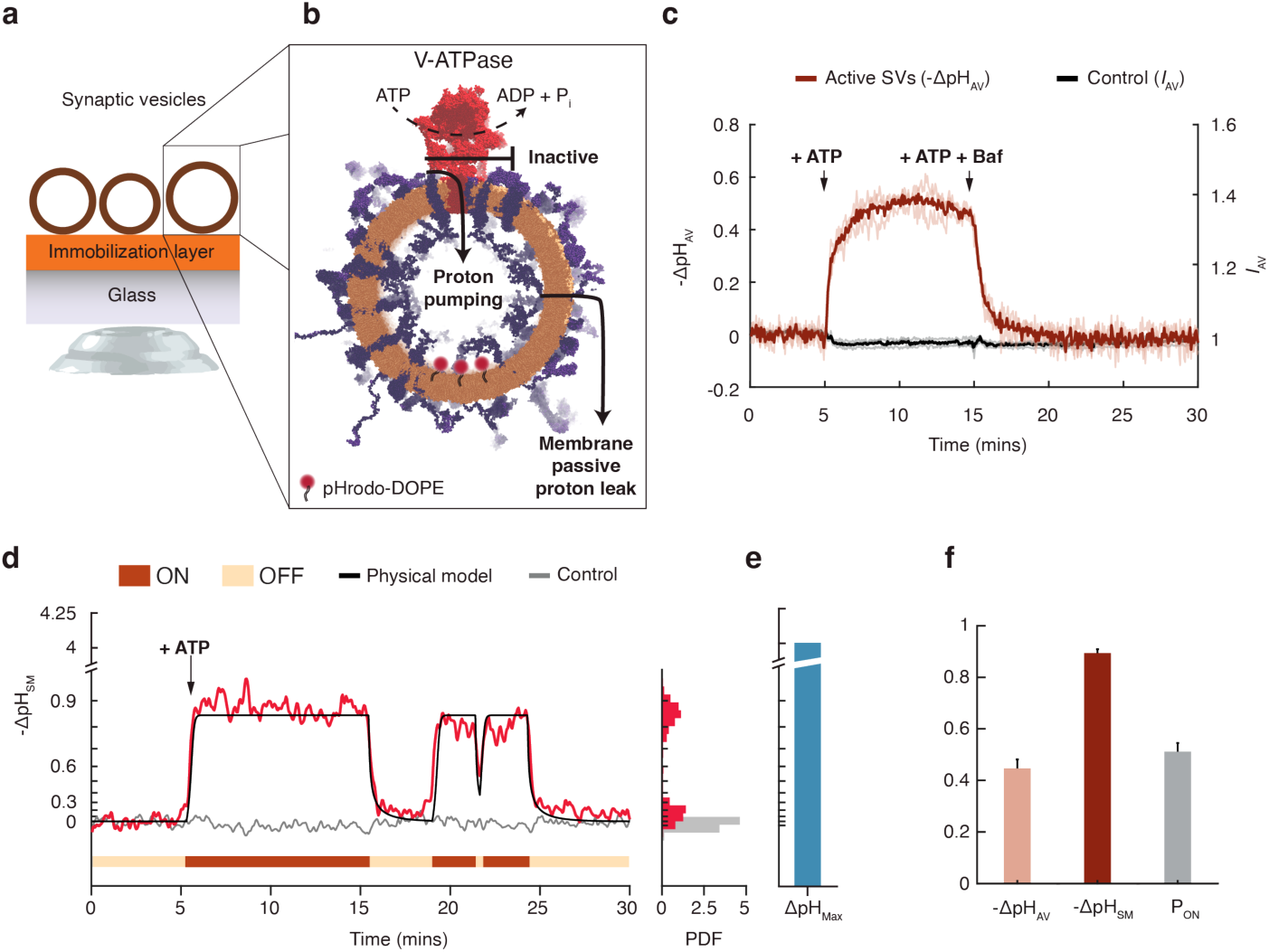
Single-molecule analysis of mammalian V-ATPase deconvolutes proton pumping from mode-switching. **a, b** Schematic of the assay used to monitor single V-ATPases acidifying the lumen of single immobilized native synaptic vesicles (SVs). **c,** Ensemble average kinetics show acidification is initiated by ATP addition and entirely blocked by the specific V-ATPase inhibitor bafilomycin A1 (Baf). Light coloured traces indicate independent replicates (n = 3); inactive vesicles are henceforth referred to as ‘control’. Conversion of fluorescence intensity (*I*_AV_) to pH was performed using a single vesicle-based calibration methodology (Supplementary Fig. 2). **d,** Representative trace of a single V-ATPase exhibiting stochastic switching between active (ON) and inactive (OFF) modes. Black line is the fit to a non-equilibrium physical model (Extended Data Fig. 1 and Supplementary Discussion). **e**, Acidification gradient (-ΔpH ∼0.5–1) is limited by passive proton leakage rather than saturation of the sensor which can report up to -ΔpH ≈ 4. **f,** Ensemble average acidification (ΔpH_AV_) is the product of single molecule acidification (ΔpH_SM_) times the probability to be in the active mode (P_ON_). Error bars denote mean ± s.e.m. (n = 5).

Single-molecule proton pumping kinetics (Fig. 1d, E.D. Fig. 1) revealed that the V-ATPase is not active continuously over time; instead, it stochastically switches between ultralong-lived but reversible active and inactive modes^8^. Since mode-switching has been observed also in a structurally distinct P-type proton pump^9^, it is likely unrelated to the irreversible dissociation of the V_1_ from the V_O_ domain^30^ but has been attributed to unfolding of small structural motifs^31^. Mode-switching remained hidden in ensemble average kinetics that were only able to capture a continuous monotonic acidification of the SV lumen (Fig. 1c). Acidification plateaus reflect a well-understood dynamic equilibrium between active proton pumping (influx into the SVs) and passive proton leakage through the lipid membrane (efflux out of the SVs). A non-equilibrium physical model^8,9,32^ (Fig. 1d black line) allowed us to deconvolve these two opposing proton kinetics, fit kinetic traces and quantitatively determine the single molecule pumping rates of the V-ATPase ∼2 H^+^s^-1^ (E.D. Fig. 1). Thus, during an active mode, whose lifetime spans timescales of minutes to hours, a V-ATPase can undergo thousands of cyclic conformational interconversions and ATP/H^+^ turnovers before reversibly transiting to an inactive mode^8^.

Mode-switching has been rigorously demonstrated for the V-ATPase and a P-type proton-pump purified and reconstituted in proteoliposomes^8,9^. In both cases, regulation can occur either through the intrinsic transport cycle or by tuning the switching probability between the active and inactive modes^8,9^. Thus, ensemble average acidification (ΔpH_AV_) is a convolution of single molecule acidification (ΔpH_SM_) and of the probability to be in the active mode (P_ON_):

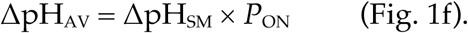

### The most potent V-ATPase inhibitors regulate mode-switching but not pumping rates

Bafilomycin A1 (hereafter bafilomycin) is a macrolide antibiotic produced by *Streptomyces griseus* (Fig. 2a) and is the most potent (IC_50_ ∼200 pM) and the most frequently used inhibitor of V-ATPase^5^. Structures of the complex show that bafilomycin binds to the membrane-embedded c-ring of the Vo domain. This interaction is hypothesized to slow down the rotation of Vo and thereby directly reduce the rate of proton transport across the membrane^5,7^. To deconvolute the molecular mechanism of inhibition, we characterised in a concentration-dependent manner the capacity of bafilomycin to inhibit the activity of single V-ATPases (Fig. 2b).

**Fig. 2.**
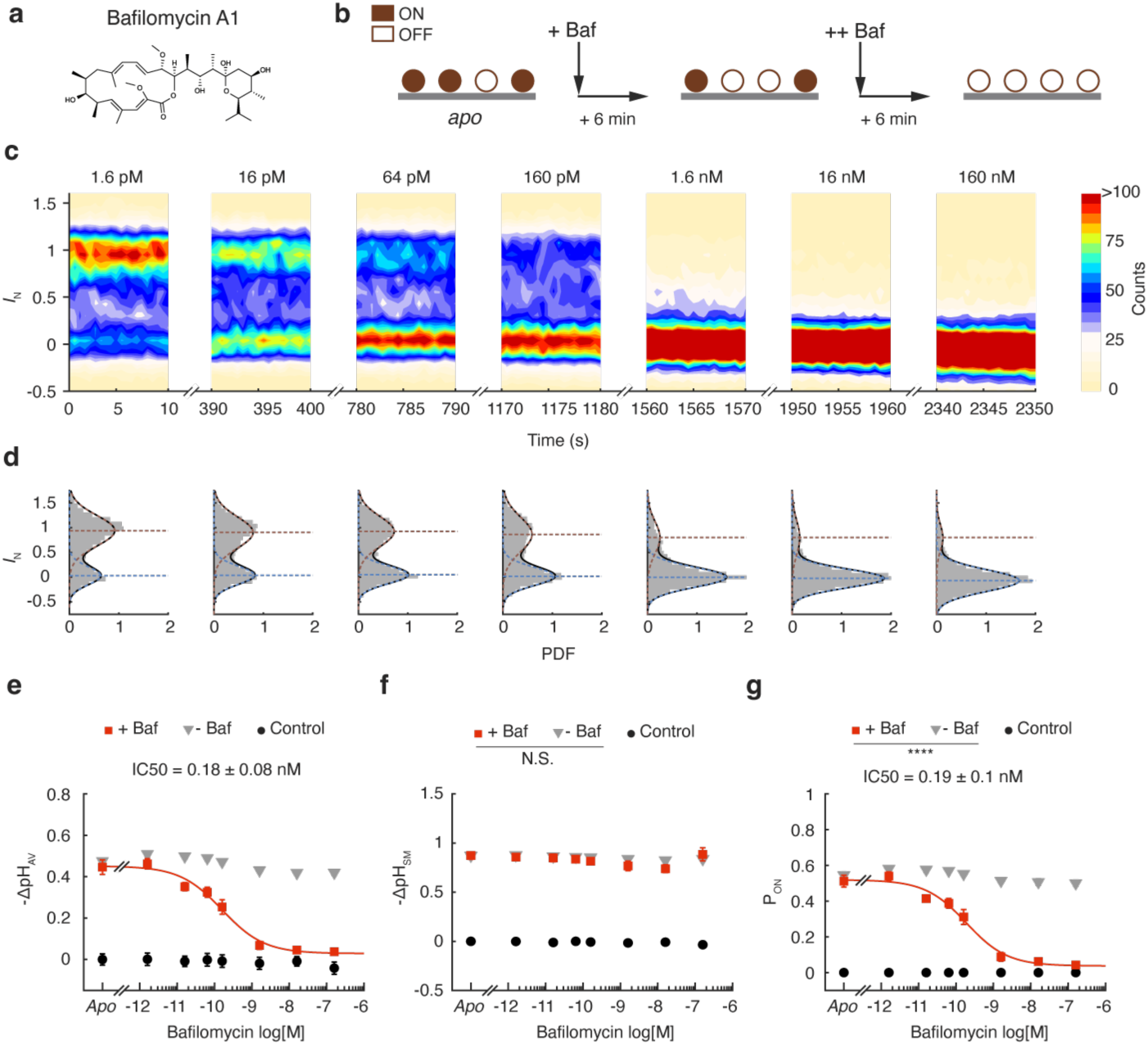
Bafilomycin inhibits V-ATPase active-mode probability but not the proton pumping of active molecules. **a,** Chemical structure of the V-ATPase inhibitor bafilomycin A1 (Baf). **b,** Schematic of in situ dose–response single-molecule imaging. **c,** Contour plots of normalized fluorescence intensity (*I*_N_) over time and Baf concentration. V-ATPases exhibit discrete transitions between inactive (*I*_N_ ≈ 0) and active (*I*_N_ ≈ 1) modes. **d,** Histograms of accumulated intensities fit to a double Gaussian distribution; increasing Baf shifts the population toward the inactive state without altering the *I*_N_ value of the active state. **e-g,** Experiments in the presence (red squares, *n* = 5, mean ± s.e.m.) and absence of Baf (grey triangles, *n* = 4, mean ± s.e.m.). Ensemble-average acidification (ΔpH_AV_) decreases with Baf in a dose-dependent manner (e). Proton pumping remains unchanged across all Baf concentrations (f); dose × condition interaction, two-way ANOVA; n.s. (P = 0.22). Active-mode probability (P_ON_) decreases with Baf in a dose-dependent manner (g); ****P < 10⁻¹⁰, dose × condition interaction, two-way ANOVA.

The stability of the sample (see Fig. 1c in reference^8^, and controls in Fig. 2e-g, grey and black markers, and E.D. Fig. 2, 3) allowed us to perform full titrations on the same single SVs and single V-ATPase molecules, Fig. 2b. We were thus able to normalise the total fluorescence intensity of individual SVs (*I*_N_) to their minimum/maximum values (-ATP and +ATP respectively), so that 0 ≤ *I*_N_ ≤ 1. This normalization corrected for the naturally occurring variability in single SV passive proton permeability and single V-ATPase pumping rates, which together determine the maximal established ΔpH_SM_ (see Supplementary Discussion). It also corrected for variabilities in single SV labelling and size, which determine the initial (-ATP) total intensity of each SV. A typical titration is shown in Fig. 2c and 2d, respectively, as population contour plots and histograms of *I*_N_. We quantified the occurrence of inactive and active V-ATPases (*I*_N_ = 0 and *I*_N_ = 1) and thus the occupancy of the two modes using a maximum likelihood estimation-based double-Gaussian fit.

Titration of bafilomycin inhibited ΔpH_AV_ in an entirely canonical manner, Fig. 2e. Strikingly, however, we did not observe any bafilomycin-induced downregulation of single molecule pumping, ΔpH_SM_, throughout the entire inhibition range, Fig. 2f (red markers), Fig. S4. Instead, titration of bafilomycin causes a pronounced shift from active to inactive modes whereby P_ON_ is reduced in a dose-dependent manner with an IC_50_ = 180 pM ± 10 pM (Fig. 2g), in excellent agreement with classical ensemble measurements^5^.

Single-molecule measurements of proton pumping thus reveal that bafilomycin does not slow down the canonical ATP-hydrolysis/rotation/proton-transport cycle of the V-ATPase. Bafilomycin regulates solely the interconversion between active and inactive modes. Consequently, the bafilomycin-induced inhibition of acidification observed in classical ensemble average experiments (Fig. 1c, 2e) can be attributed entirely to the regulation of mode-switching.

To understand whether pharmacological inhibition mediated via mode-switching is a unique property of bafilomycin, we tested two more widely employed inhibitors. Concanamycin A (hereafter concanamycin) is a macrolide antibiotic related to bafilomycin, which, despite its different side groups and stereochemistry, is believed to share a similar inhibitory mechanism^7^ (Fig. 3a). Its binding site to the V-ATPase has not been structurally characterised. However, structural models suggest the site is shared with bafilomycin, albeit concanamycin being larger and having more contact points with the V-ATPase^7^. Diphyllin is a plant-derived lignan lactone with a rigid tricyclic structure that is structurally unrelated to bafilomycin (Fig. 4a) and whose binding site for the V-ATPase is unknown^34^. Diphyllin and its derivatives, designed to increase bioavailability, have been investigated for use in anticancer, antiviral, and lysosome-targeting therapies^35^.

**Fig. 3.**
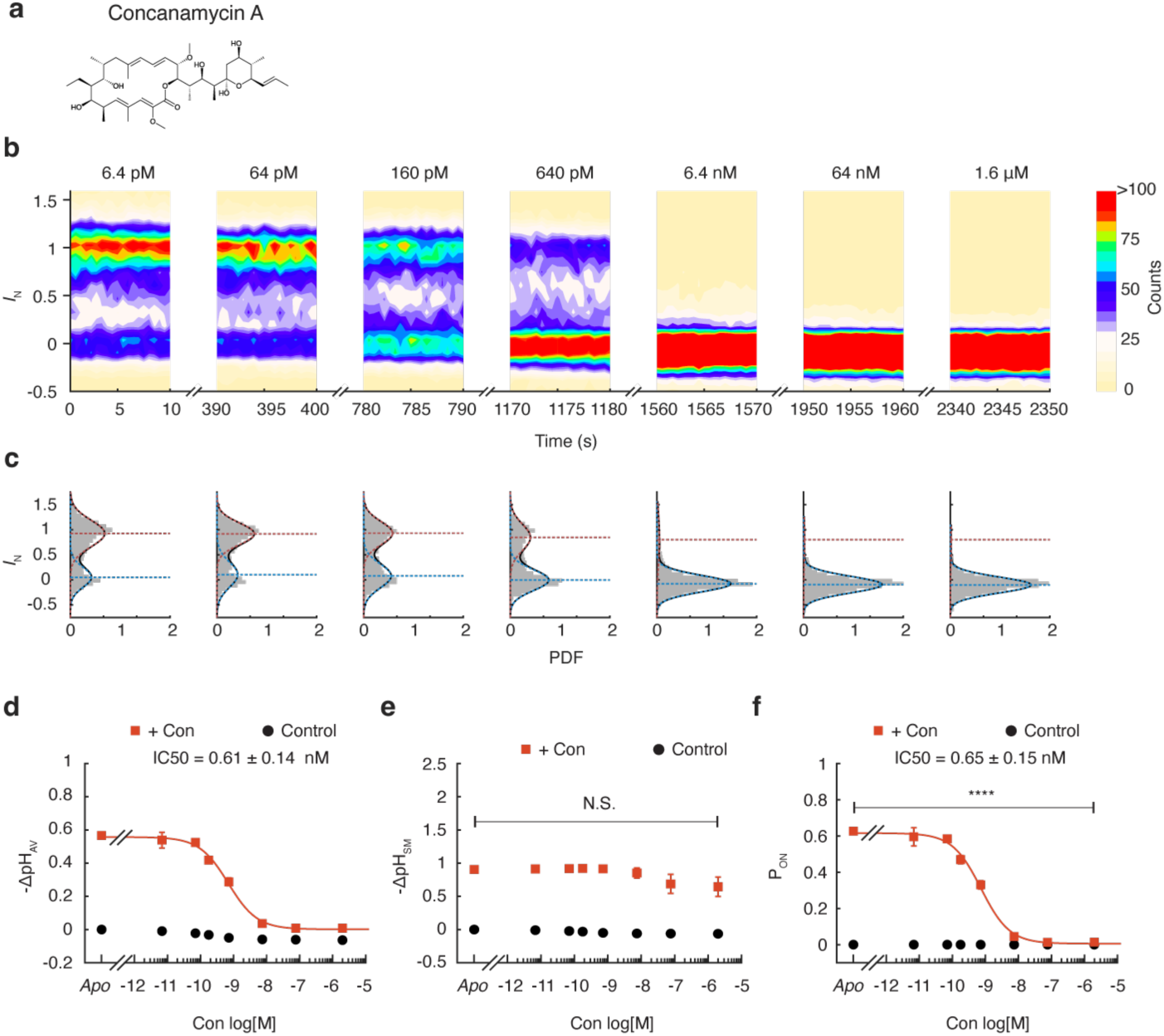
Concanamycin inhibits V-ATPase active-mode probability but not the proton pumping of active molecules. **a,** Chemical structure of concanamycin A (Con). **b,** Contour plots of normalized fluorescence intensity over time reveal discrete transitions between inactive (*I*_N_ ≈ 0) and active (*I*_N_ ≈ 1) modes across Con concentrations. **c,** Histograms of accumulated intensities fit to a double Gaussian distribution; increasing Con shifts occupancy toward the inactive mode without altering active mode intensity. **d,** Ensemble-average acidification (ΔpH_AV_) decreases with Con in a dose-dependent manner, following a classical isotherm; *n* = 3, mean ± s.e.m. **e,** Single-molecule acidification (ΔpH_SM_) remains unchanged across inhibitor concentrations; *n* = 3, mean ± s.e.m., one-way ANOVA, n.s. (P = 0.14). **f,** Active-mode probability (P_ON_) decreases with increasing Con; *n* = 3, mean ± s.e.m., P < 10⁻¹⁰, one-way ANOVA.

**Fig. 4.**
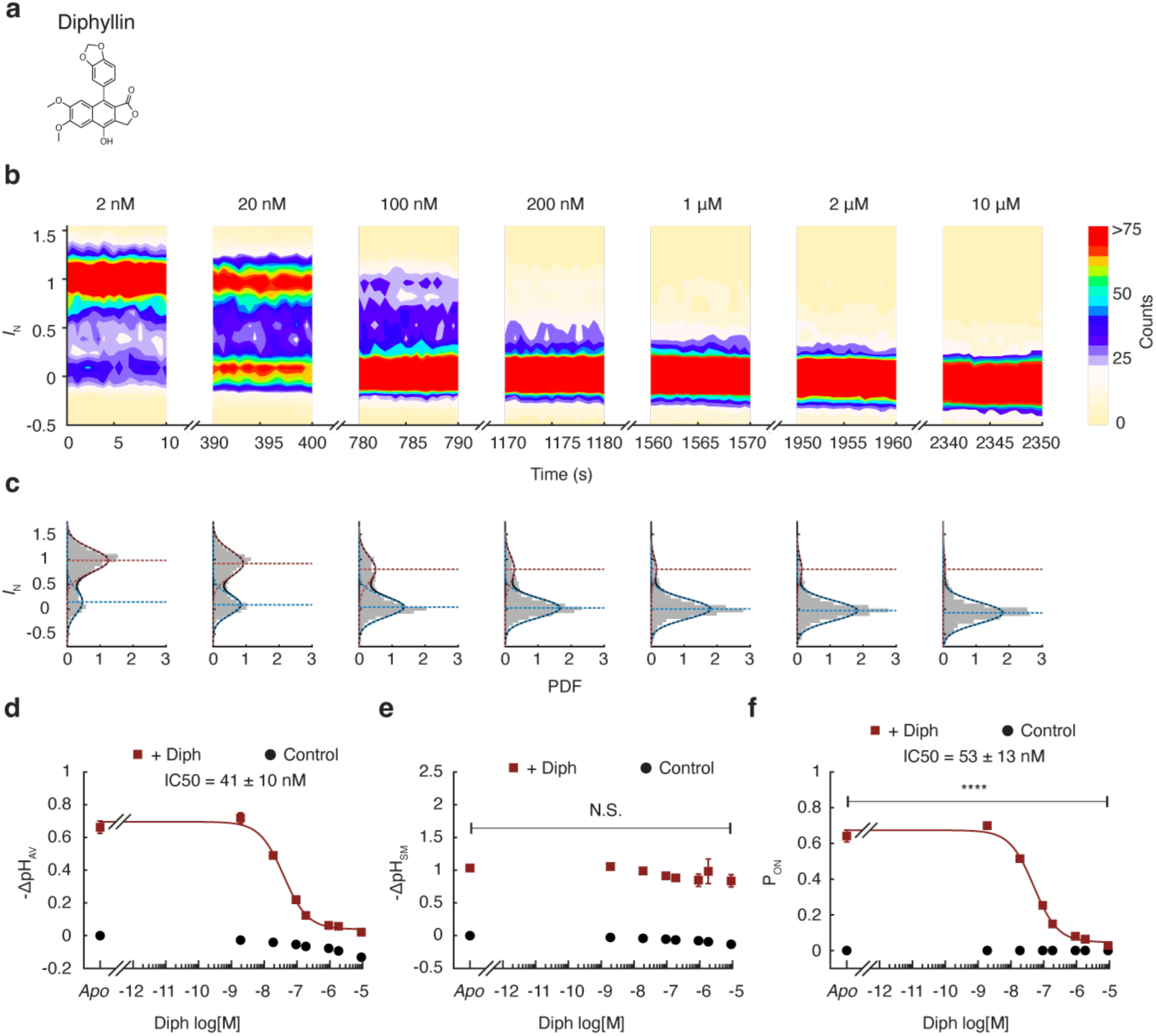
Diphyllin inhibits V-ATPase active-mode probability but not the proton pumping of active molecules. **a,** Chemical structure of diphyllin (Diph). **b,** Contour plots of normalized fluorescence intensity over time reveal discrete transitions between inactive (*I*_N_ ≈ 0) and active (*I*_N_ ≈ 1) modes across Diph concentrations. **c,** Histograms of accumulated intensities fit to a double Gaussian distribution; increasing Diph shifts occupancy toward the inactive mode without altering active mode intensity. **d,** Ensemble-average acidification (ΔpH_AV_) decreases with Diph in a dose-dependent manner, following a classical isotherm; *n* = 3, mean ± s.e.m. **e,** Single-molecule acidification (ΔpH_SM_) remains unchanged across inhibitor concentrations; *n* = 3, mean ± s.e.m., one-way ANOVA, n.s. (P = 0.57). **f,** Active-mode probability (P_ON_) decreases with increasing Diph; *n* = 3, mean ± s.e.m., P < 10⁻¹⁰, one-way ANOVA.

The ensemble average IC_50_ values we determined for concanamycin and diphyllin (610 pM and 41 nM, Fig. 3d, 4d) matched closely published values^7,34^. However, the single-molecule dose-response measurements confirmed that neither compound was able to slow down the intrinsic proton pumping rates in a statistically significant manner (Fig. 3e, 4e, S4). Notably, both concanamycin and diphyllin shared the same molecular mechanism of V-ATPase inhibition as bafilomycin, namely through modulation of the probability to exit the canonical transport cycle and enter a long-lived inactive mode (Fig. 3f, 4f).

### Mode-specific and differentially efficient regulation of mode lifetimes

To further elucidate how inhibitors regulate the active mode probability, *P*_on_, we analysed all individual mode switching events in thirty-minute-long kinetic traces of proton pumping by single V-ATPases. This allowed us to deconvolve the regulatory effects of inhibitors on the lifetimes of the active and inactive modes (*τ*_ON_ and *τ*_OFF_).

As shown in Fig. 5a-c, we first recorded five minutes of baseline (-ATP), then ten minutes of activity (+ATP, *apo*), followed by fifteen minutes of inhibition (+ATP, +inhibitor) at concentrations close to the IC_50_ of the compounds. We determined the time points at which the active and inactive modes stochastically start and end with single-frame precision, utilizing a previously developed Bayesian change-point-detection algorithm (Supplementary Discussion, E.D. Fig. 4) that identifies significant deviations from the baseline noise of each SV^8^. Event detection enabled us to generate histograms of event acidification (ΔpH_SM_) and event lifetimes in the *apo* condition, as well as in the presence of inhibitors (E.D. Fig. 6 and E.D. Fig. 8).

**Fig. 5.**
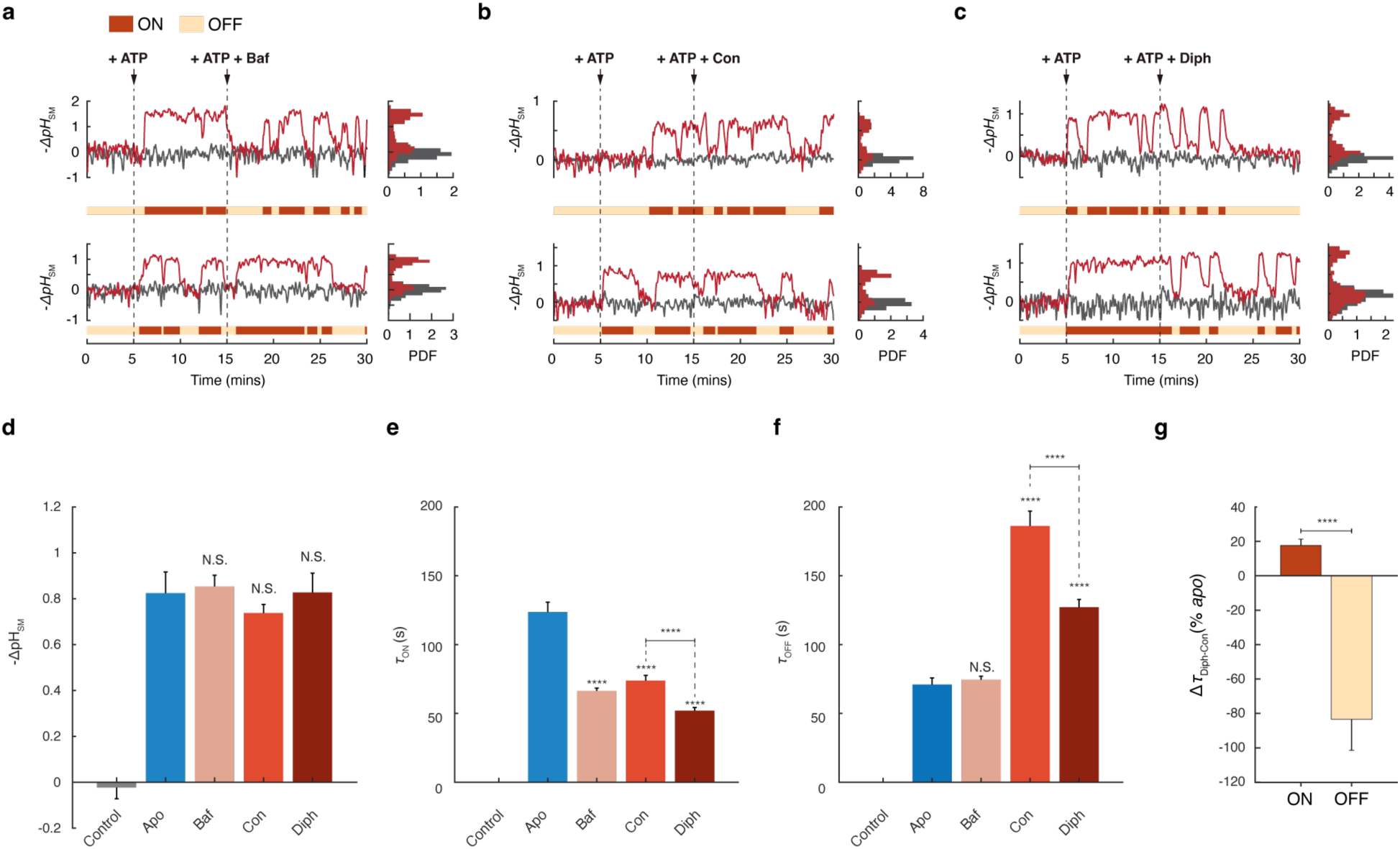
Single-molecule kinetics reveal V-ATPase inhibitors can specifically target and differentially regulate mode lifetimes. **a-c**, Typical traces of acidification kinetics of single active SVs (red) exhibit stochastic transitions between active and inactive modes. Time traces of inactive SVs are shown as a control (grey). Histograms show the distribution of pH gradients over time. Inhibitor concentrations were: 50 pM (Baf), 320 pM (Con), and 50 nM (Diph). **d**, Proton gradients (ΔpH_SM_) in the absence/presence of inhibitors. n.s. (*P* = 0.52, ± Baf); n.s. (*P* = 0.58, ± Con); n.s. (*P* = 0.72, ± Diph); two-sided two-sample Kolmogorov-Smirnov test; mean ± s.d. from independent experiments; *n* = 11 (*apo*); *n* = 3 (Baf); *n* = 4 (Con, Diph). **e,** Active mode dwell times (*τ*_ON_) in the presence and absence (*apo*) of inhibitors. \*\*\*\**P* < 10^-10^ (Baf-*apo*); \*\*\*\**P* < 10^-10^ (Con-*apo*); \*\*\*\**P* < 10^-10^ (Diph-*apo*); **** *P* = 8·10^-8^ (Con-Diph); pairwise two-sided Kolmogorov-Smirnov test performed on single vesicle data. **f**, Inactive mode dwell times (*τ*_OFF_) in the presence and absence (*apo*) of inhibitors. Bafilomycin had no apparent effect on the inactive mode dwell times. Con and Diph upregulated *τ*_OFF_. n.s. (*P* = 0.63, Baf-*apo*); \*\*\*\**P* < 10^-10^ (Con-*apo*); \*\*\*\**P* < 10^-10^ (Diph-*apo*); \*\*\*\**P* = 2·10^-5^ (Con-Diph); pairwise two-sided Kolmogorov-Smirnov test. *τ*_ON_ and *τ*_OFF_ were calculated by fitting single exponentials to the dwell time histograms. *Apo*: error bars indicate the s.d. of the mean across fits from independent experiments. *n* = 3. Inhibitors: error bars correspond to the standard error of the fit. **g,** Difference of characteristic lifetimes between Diph and Con normalized to their relative change to the *apo* condition; \*\*\*\**P =* 4·10^-8^; two-sample two-tailed z-test; error bars correspond to the standard error of the fit; *n* = 4.

**Fig. 6.**
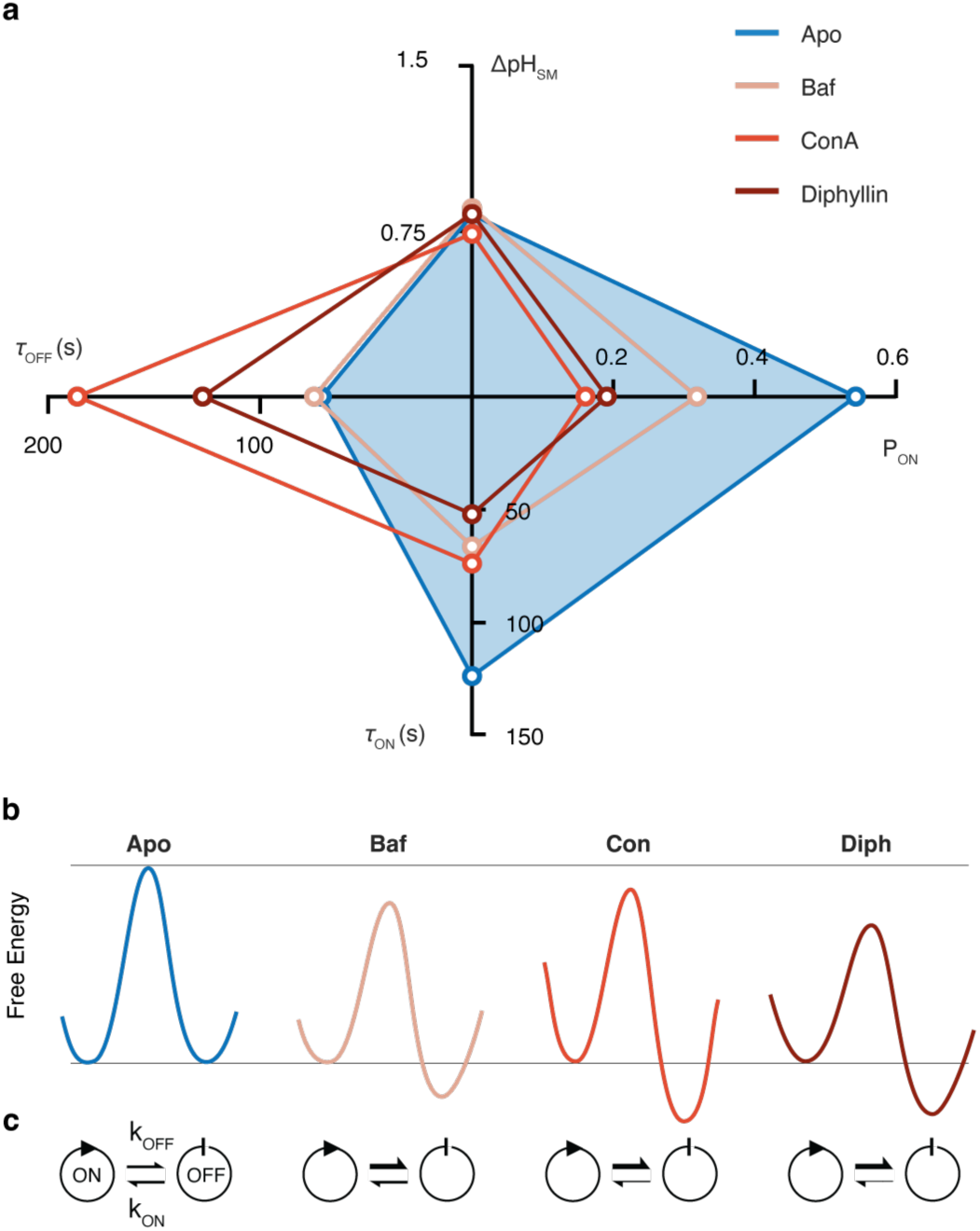
Transport inhibition through mode-switching instead of the transport cycle. **a**, Spider chart of single-molecule properties of the V-ATPase for *apo*, bafilomycin, concanamycin, and diphyllin summarises the observed compound-specific molecular mechanisms of mode-based inhibition. **b**, Illustration of free energy landscapes of mode-switching transitions in the absence and presence of inhibitors. The diagrams are informed by the data summarised in E.D. Table 1 but exaggerate the magnitude of free energy changes for visual clarity, see also Supplementary Discussion. Inhibitors can differentially reshape energy barriers and/or relative free energy minima. **c**, Mode-switching rate constants (k_ON_ and k_OFF_) reveal distinct effects of inhibitors on the interconversion between active and inactive modes. These results delineate a novel mechanism of pharmacological inhibition and a fundamentally new principle for drug design.

None of the three inhibitors reduces event-specific ΔpH_SM_ (Fig. 5d, E.D. Fig. 6), consistent with the single-molecule population average values shown in Figs. 2-4. However, analysis of mode lifetime distributions demonstrated that each inhibitor elicited a unique kinetic regulation profile (Fig. 5e, f, and E.D. Fig. 8). Specifically, bafilomycin reduced *τ*_ON_ by 50% but had no distinguishable effect on *τ*_OFF_, demonstrating that inhibitors can exhibit essentially perfect mode-specificity or "mode-bias". In contrast, concanamycin and diphyllin regulated the lifetimes of both modes, however, in an opposite manner, where they reduced *τ*_ON_, but increased *τ*_OFF,_ i.e., they destabilise the active mode but stabilise the inactive one, Fig. 5e, f. Furthermore, concanamycin and diphyllin exhibited different relative efficiencies in regulating the lifetimes of the two modes, e.g., compared to concanamycin, diphyllin was four-fold better at modulating *τ*_OFF_ versus *τ*_ON_, Fig. 5g.

To provide further perspective on the regulation of mode-switching, we examined the active-mode lifetime distributions. Specifically, we sought to distinguish whether inhibitors act by directly truncating active-mode durations through transient/reversible blockage, or by allosterically modulating the intrinsic transitions between active and inactive modes. Allosteric modulation would transform the monoexponential distribution of lifetimes into a double (or stretched) exponential^36^. Consistent with this model, Bayesian analysis (S. Table 1) revealed that bafilomycin introduces a clear second exponential component to the active-mode lifetime distribution. Interestingly, concanamycin and diphyllin did not, a result that is consistent with direct blockage^37^. These findings suggest that different inhibitors can leverage distinct mechanistic strategies to regulate V-ATPase mode-switching. The corresponding effects on the energy landscape are summarized in E.D. Table 1 and Figure 6.

## Conclusion

Our single-molecule investigation reveals a fundamentally new understanding of V-ATPase pharmacological inhibition. Contrary to the popular implicit assumption that potent inhibitors suppress proton transport by directly slowing the catalytic transport cycle, we demonstrate that the proton pumping rate of active enzymes can remain remarkably unaffected. Instead, inhibition here arises exclusively through regulation of mode-switching kinetics and by shifting occupancy from active to inactive modes. Tested inhibitors exhibited a distinct kinetic "fingerprint," selectively modulating the lifetimes of active and inactive modes in a specific and differential manner.

The mechanistic shift from catalytic suppression to mode-based pharmacological inhibition has several important implications. First, it resolves the apparent paradox wherein ensemble-averaged V-ATPase measurements indicate strong inhibition despite unaltered single-molecule proton pumping catalytic rates. Second, it identifies tool compounds to isolate and investigate the physiological roles of mode-switching^8^. Indeed, previous studies employing bafilomycin to perturb synaptic vesicle electrochemical gradients were, in effect, observing how altering the duration and frequency of stochastic, all-or-none mode transitions impacts neurotransmission^38–40^. Third, this insight significantly expands the conceptual framework of transporter pharmacology. Given that mode-switching has been documented across diverse primary^8,9^ and secondary^10–13^ active transporters, our findings offer proof of principle for innovative drug-design strategies aimed explicitly at modulating mode occupancy and switching kinetics, rather than directly targeting catalytic turnover and the substrate transport cycle.

## Methods

### Chemicals

Phospholipids were procured from Avanti Polar Lipids Inc., while chemicals for buffers, detergents, and other reagents were obtained from Sigma-Aldrich, unless specified differently. Bafilomycin was purchased from Sigma-Aldrich and MedChemExpress. Concanamycin A and diphyllin were purchased from MedChemExpress.

### The lipid-conjugated *pH* sensor PE-pHrodo

1,2-dioleoyl-sn-glycero-3-phosphoethanolamine (DOPE) lipids were linked to pHrodo™ Red, succinimidyl ester (Invitrogen). The synthesis, purification, and characterization of the lipid-conjugated pH sensor have been described in detail^41^.

### Synaptic vesicles from rat brain

Synaptic vesicles (SVs) were extracted from rat brain following previously published methods^21^. In summary, an SV-enriched fraction, denoted as LP2, was generated through differential centrifugation and subsequently subjected to continuous sucrose density gradient centrifugation. The portion located between 0.04 and 0.4 M sucrose (highlighted in color) was collected and further separated through chromatography using controlled-pore glass beads (CPG). Following size exclusion chromatography, the SVs were sedimented via ultracentrifugation, resuspended in a solution containing 320 mM sucrose and 5 mM HEPES at pH 7.4, and promptly frozen for storage at -80 °C until needed. For more detailed information, please refer to Supplementary Fig. 1. The protein concentration in this purified SV fraction ranges from 1.5 mg/ml to 2.5 mg/ml.

This purified SV fraction is a mixture of all SVs found in the whole brain. Recently, the distribution of all known vesicular neurotransmitter transporters has been quantified within this preparation^42^. Consequently, nearly 80% of SVs are glutamatergic, and 15% are GABAergic, which confirms findings from a prior study^28^. Vesicles containing other transporters (e.g., for dopamine, acetylcholine, etc.) constitute only a small fraction, except for the Zn2+ transporter ZnT3, which co-localizes with approximately one-third of the glutamatergic vesicles^43^.

Adult Wistar rats were procured from Charles River Laboratories or Janvier and were housed until they reached 5 to 6 weeks of age under a 12:12-hour light/dark cycle, with access to food and water ad libitum. TierSchG (Tierschutzgesetz der Bundesrepublik Deutschland, Animal Welfare Law of the Federal Republic of Germany) as documented by 33.23-42508-066-§11, dated Nov 16^th^, 2023 ("Erlaubnis zum Halten von Wirbeltieren zur Versuchszwecken", "Permission to keep vertebrates for experimental purposes") by the Niedersächsisches Landesamt für Verbraucherschutz und Lebensmittelsicherheit (Lower Saxony State Office for Consumer Protection and Food Safety).

### Surface preparation and immobilization of synaptic vesicles

Surface functionalization followed established protocols^23–26^. Briefly, glass slides (0.17 ± 0.01 mm thickness) were cleaned by sequential sonication in 2% (v/v) Hellmanex III, MilliQ water, 99% (v/v) ethanol, and 99% (v/v) methanol. Slides were dried under nitrogen and plasma-etched for 3 minutes to remove residual contaminants. Cleaned slides were affixed to Ibidi stickySlide VI 0.4 chambers.

Surfaces were functionalized by incubating a 1 mg mL⁻¹ solution of PLL-g-PEG and PLL-g-PEG-biotin (100:1 ratio; SuSoS) for 30 minutes, followed by rinsing with 15 mM HEPES. Neutravidin (0.1 mg mL⁻¹ in 15 mM HEPES) was then added for 10 minutes and washed out with activity buffer (300 mM glycine, 2 mM MOPS, 2 mM MgSO₄, pH 7.1–7.15 at ∼23 °C, adjusted with Tris).

After equilibration with activity buffer, synaptic vesicles (SVs) suspended in the same buffer were introduced and incubated until ∼1500–2000 vesicles were immobilized within an 81.92 µm × 81.92 µm field of view (FOV). Unbound vesicles were subsequently flushed out.

### Lipid labelling of intact synaptic vesicles

To embed fluorescent and biotinylated lipids into native synaptic-vesicle (SV) membranes, 10 µL pHrodo-DOPE (1 mg mL⁻¹) and 5 µL DSPE-PEG(2000)-biotin (10 mg mL⁻¹) were dissolved in 85 µL chloroform. The solvent was evaporated under nitrogen to form a thin lipid film, and then the sample was dried under vacuum for at least 30 minutes.

The film was rehydrated with 20 µL purified SV suspension diluted in 80 µL activity buffer, mixing gently by repeated pipetting until fully dissolved. The suspension was agitated at 600 rpm for 10 min (room temperature) and centrifuged at 13 000 rpm for 1 min to pellet debris and dye aggregates while leaving the SVs in the supernatant. The pellet was subsequently discarded and only the supernatant was kept. SVs were then aliquoted (10 µL) and subjected to one freeze–thaw cycle before use.

### Image acquisition

Total internal reflection fluorescence (TIRF) microscopy was performed using a commercial Olympus TIRF system (Olympus Europa S.E. & Co. KG, Hamburg, Germany) equipped with a 532 nm solid-state laser (Olympus Cell*) and a UApoN ×100/1.49 NA oil immersion objective. Emission was collected through a 582.5/75 bandpass filter and separated from excitation via a 585/75 beam splitter. A 532/10 excitation filter was used to ensure selective fluorophore excitation. Images were acquired using an iXon 897 EMCCD camera (Andor Technology, Belfast, UK). Focus was stabilized using the Olympus Zero Drift Correction (ZDC) module.

Image acquisition and analysis were performed using Olympus cellSens software (v3.2). During kinetic measurements, samples were illuminated for 500 ms per frame, with a 3 s cycle time. The XY stage cycled between four fields of view (FOVs), and the laser remained off during stage movement. For snapshot experiments, 20-frame stacks were recorded per condition, cycling through six FOVs between stacks.

---

**Extended Data Fig. 1.**
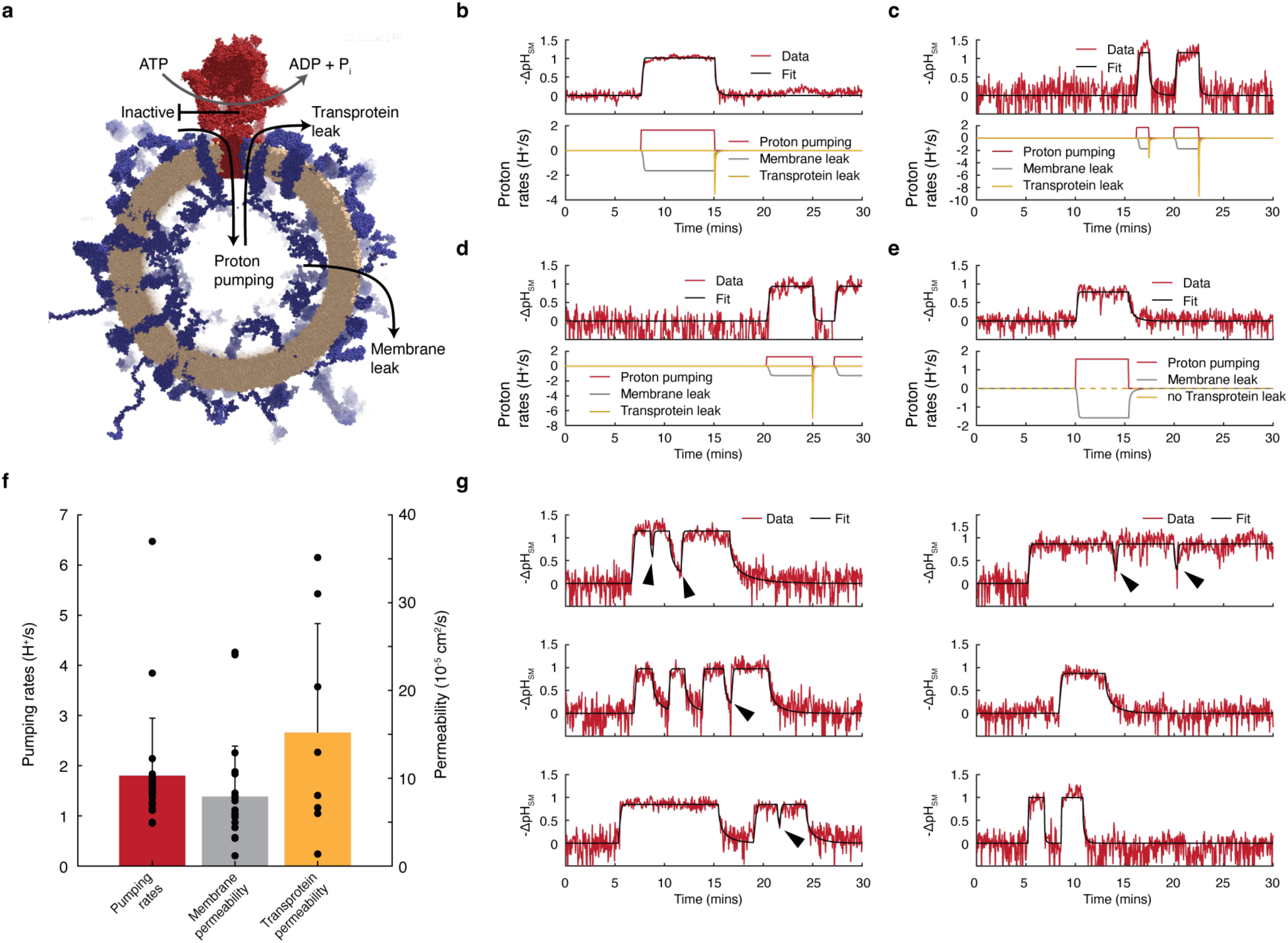
Non-equilibrium modelling reveals single-molecule proton transport kinetics and leakage pathways. **a**, Schematic of model parameters. **b–e**, Top: Representative single-molecule traces showing stochastic mode-switching dynamics. Active proton pumping is intermittently interrupted by inactive and proton-leaky modes. During active phases, a dynamic equilibrium is established between pumping and leakage, resulting in a single acidification plateau. During an active mode, which can last up to hours, a V-ATPase may complete thousands of conformational cycles and ATP/H⁺ turnover events before randomly switching to an inactive modes. Leakage arises from both membrane diffusion and transprotein efflux. Transprotein leakage initiates immediately upon V-ATPase deactivation and displays distinct temporal kinetics. Traces were fitted using a non-equilibrium model (see Supplementary discussion), allowing for the independent estimation of pumping rates and permeabilities. Bottom: Modelled rates for proton pumping (red), membrane leak (grey), and transprotein leak (yellow). In b–d, efflux is dominated by transprotein leakage; in e, by membrane diffusion. **f**, Model-derived proton transport rates. Pumping rates (red) average 1.8 ± 1.1 H⁺ s⁻¹. Membrane (grey) and transprotein (yellow) permeabilities are estimated at 8 ± 5 × 10⁻⁵ cm s⁻¹ and 15 ± 12 × 10⁻⁵ cm s⁻¹, respectively. Error bars, s.d.; n = 24 (pumping), n = 24 (membrane), n = 8 (transprotein). **g**, Additional single-molecule traces with model fits. Arrows indicate mode-switching events where equilibrium was not reached (ΔpH_SM_ ≈ max or 0). Transprotein leakage modes were previously reported⁸,⁹ but are not further analysed here.

**Extended Data Fig. 2.**
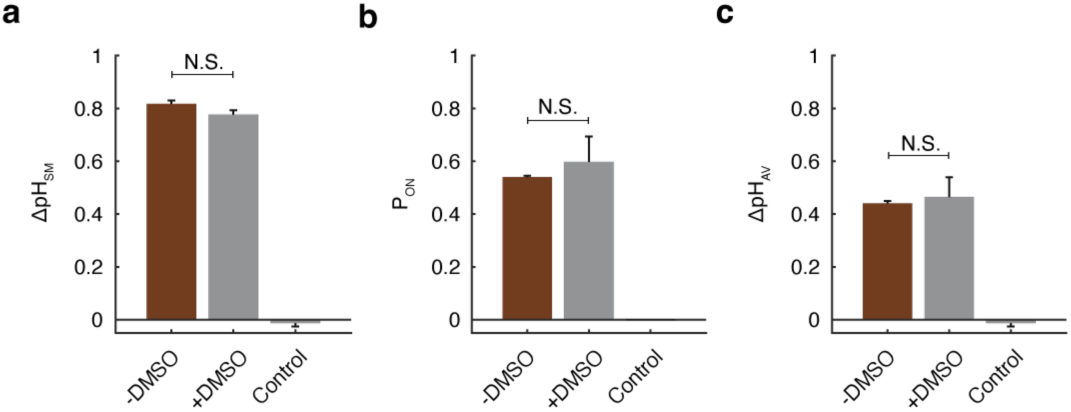
DMSO does not affect single-molecule and ensemble average pH gradients and active mode probabilities. **a**, Single-molecule pH gradients under apo conditions (+ATP only) with and without DMSO show no significant difference. Error bars, s.e.m.; n = 4; n.s. (P = 0.18); two-sided Student’s t-test. **b**, Active mode probabilities remain unchanged in the presence of DMSO under apo conditions. Error bars, s.e.m.; n = 4; n.s. (P = 0.63); two-sided Student’s t-test. **c**, Ensemble average acidification is also unaffected by DMSO. Error bars, s.e.m.; n = 4; n.s. (P = 0.18); two-sided Student’s t-test.

**Extended Data Fig. 3.**
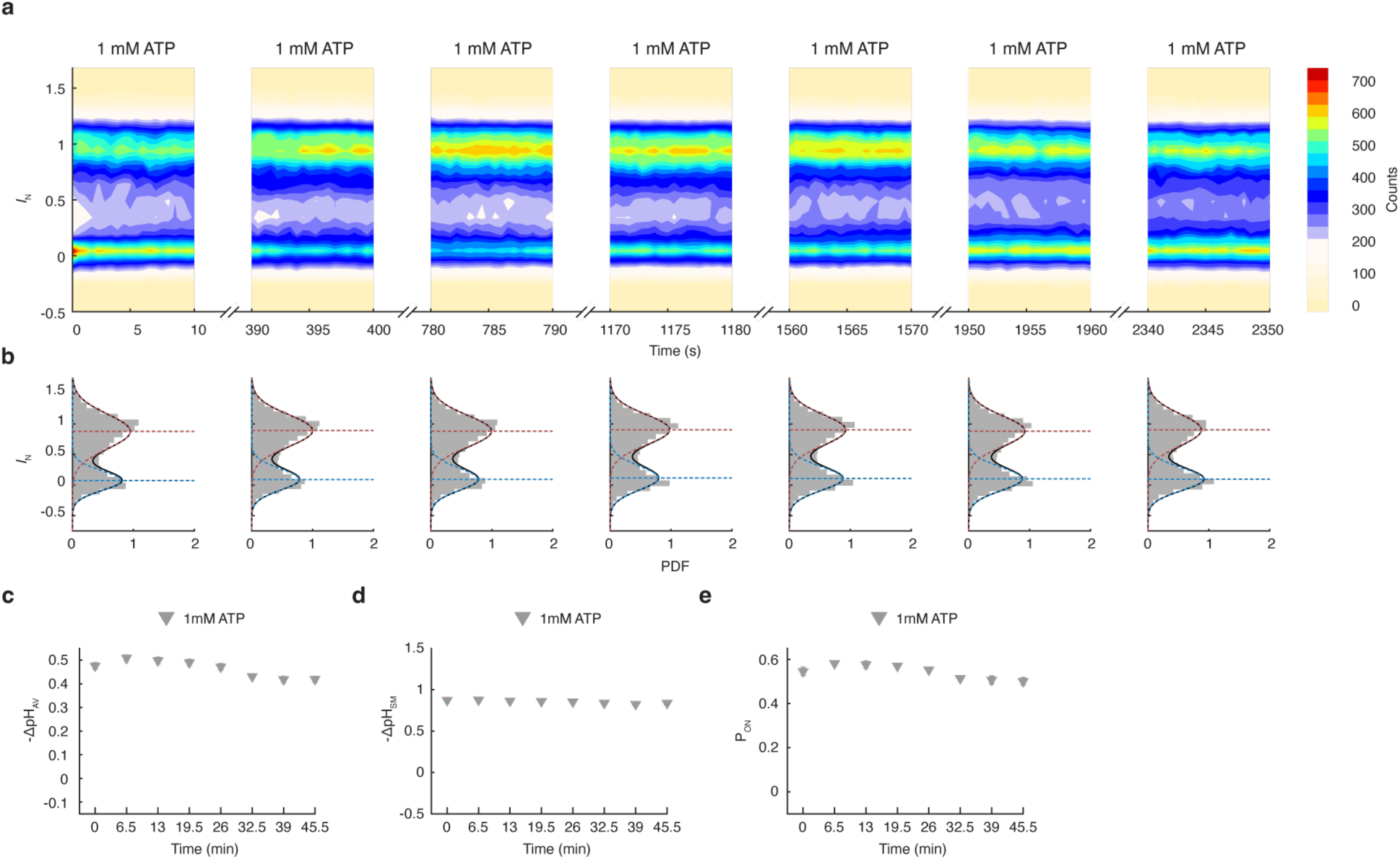
Control confirms the stability of ΔpH_SM_ over periods of 45 minutes. **a,** Contour plots of normalized fluorescence intensity over time reveal discrete transitions between inactive (*I*_N_ ≈ 0) and active (*I*_N_ ≈ 1) modes at a constant ATP concentration (1 mM ATP). **b,** Histograms of accumulated intensities fit to a double Gaussian distribution; mode occupancies of active and inactive modes remain constant over time. **c,** Ensemble-average acidification (ΔpH_AV_) remains constant over time; *n* = 4, mean ± s.e.m. **d,** Single-molecule acidification (ΔpH_SM_) remains unchanged across inhibitor concentrations; *n* = 4, mean ± s.e.m., one-way ANOVA, n.s. (P = 0.57). **e,** Active-mode probability (P_ON_) remains constant over time; mean ± s.e.m. from independent experiments; *n* = 4.

**Extended Data Fig. 4.**
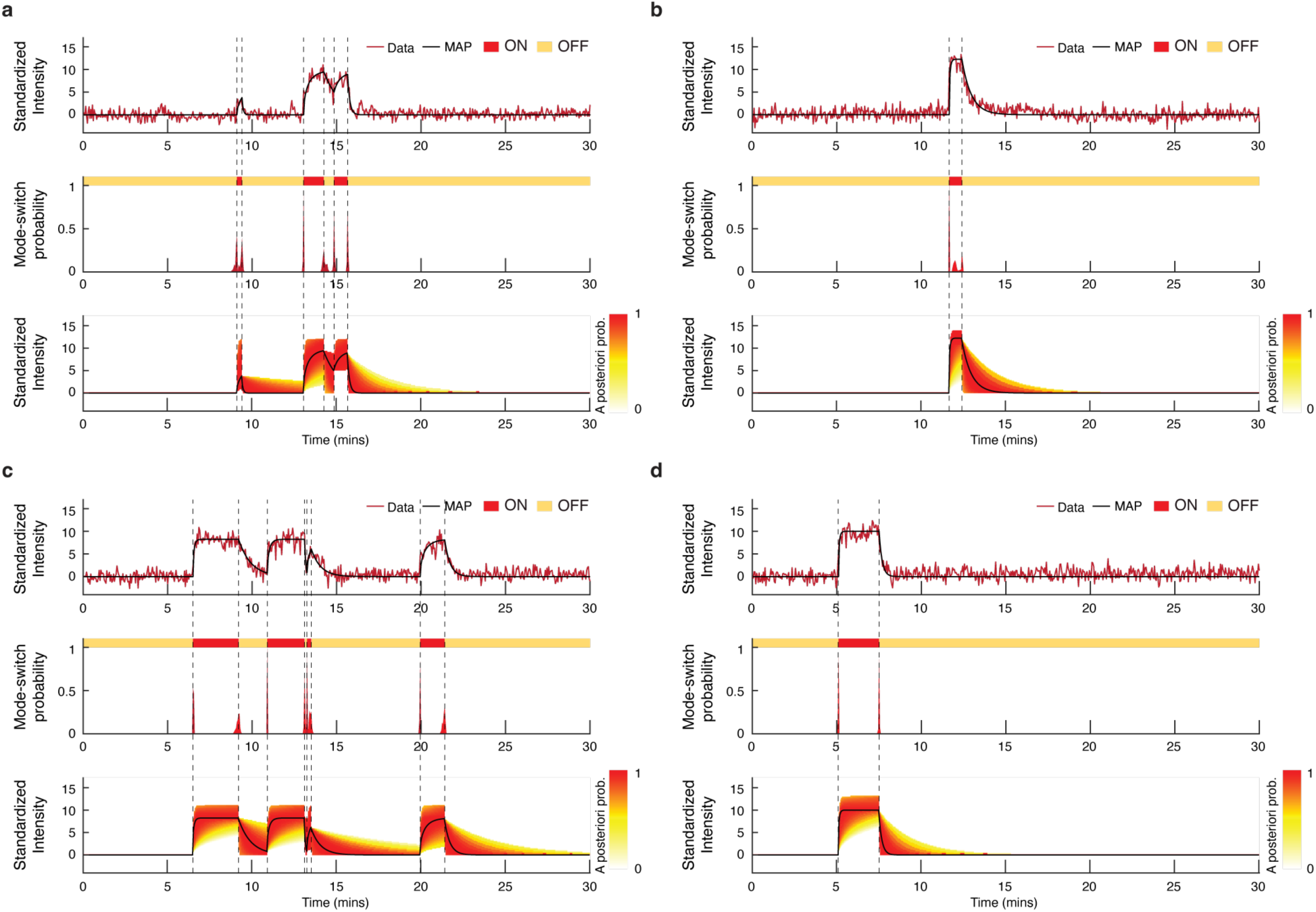
MAP-based detection of mode-switching events using a stochastic exponential model. **a–d**, Single-molecule fluorescence traces were analysed using Bayesian stochastic filtering based on a piecewise exponential model of V-ATPase activity. The signal reflects alternating proton-pumping and inactive states, modelled by exponential increases or decays in fluorescence intensity. Change-points (mode-switching events) were identified from the posterior distribution of the signal, with events marked at time points reaching ≥99% confidence (dashed lines). MAP (maximum a posteriori) estimates were derived by optimizing over the joint posterior distribution of model parameters and change-point times (see Supplementary Methods and reference^8^). Red curves in the probability panels indicate the marginal posterior distribution for switching events. Heatmaps show the posterior probability of signal location; MAP estimates are overlaid in black.

**Extended Data Fig. 5.**
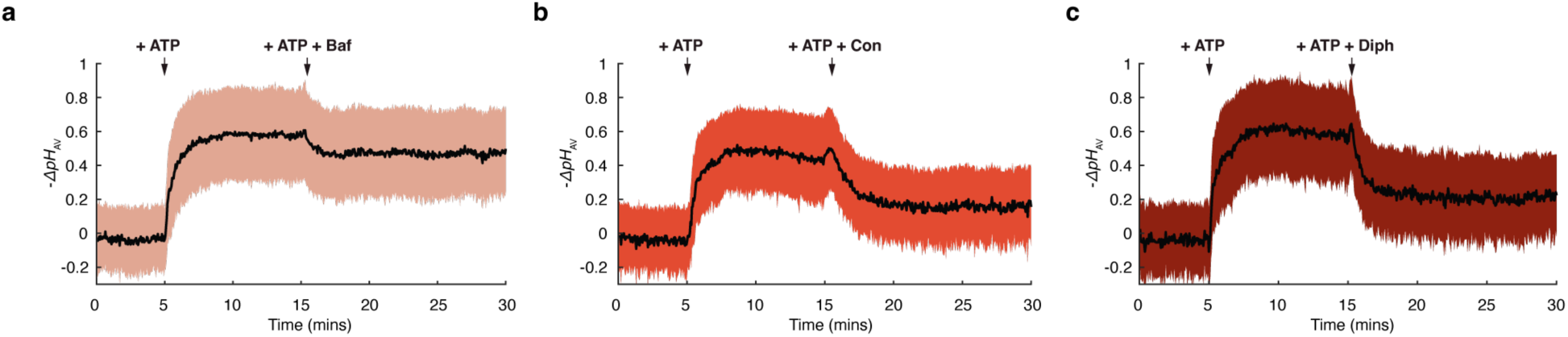
Partial reduction of ensemble average acidification by the V-ATPase at intermediate concentration of inhibitors. **a-c**, Ensemble average acidification kinetics of 834, 519, and 452 single, active SVs in the presence of bafilomycin, concanamycin, and diphyllin, respectively. Activity is initiated upon the addition of ATP. After 10 minutes, inhibitors are injected, and net acidification is notably reduced. Black line and coloured areas reflect the mean and one s.d. between SVs. The ensemble average trace for Baf has been corrected for bleaching.

**Extended Data Fig. 6.**
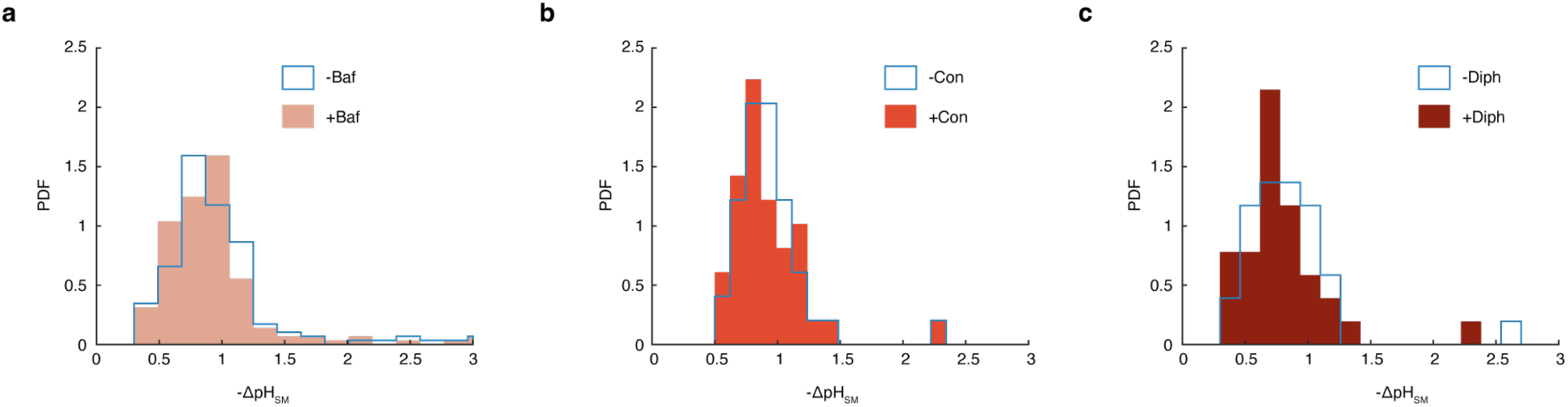
Inhibitors do not affect the levels of acidification established by single-V-ATPases. **a-c**, Distributions of single-molecule acidification levels in SVs by the V-ATPase in the presence and absence of inhibitors estimated from acidification plateaus of single-molecule kinetic measurements. Concentrations of inhibitors used were: 50 pM Baf; 320 pM Con; 50 nM Diph. n.s. (*P* = 0.52, ± Baf); n.s. (*P* = 0.58, ± Con); n.s. (*P* = 0.72, ± Diph); two-sided two-sample Kolmogorov-Smirnov test.

**Extended Data Fig. 7.**
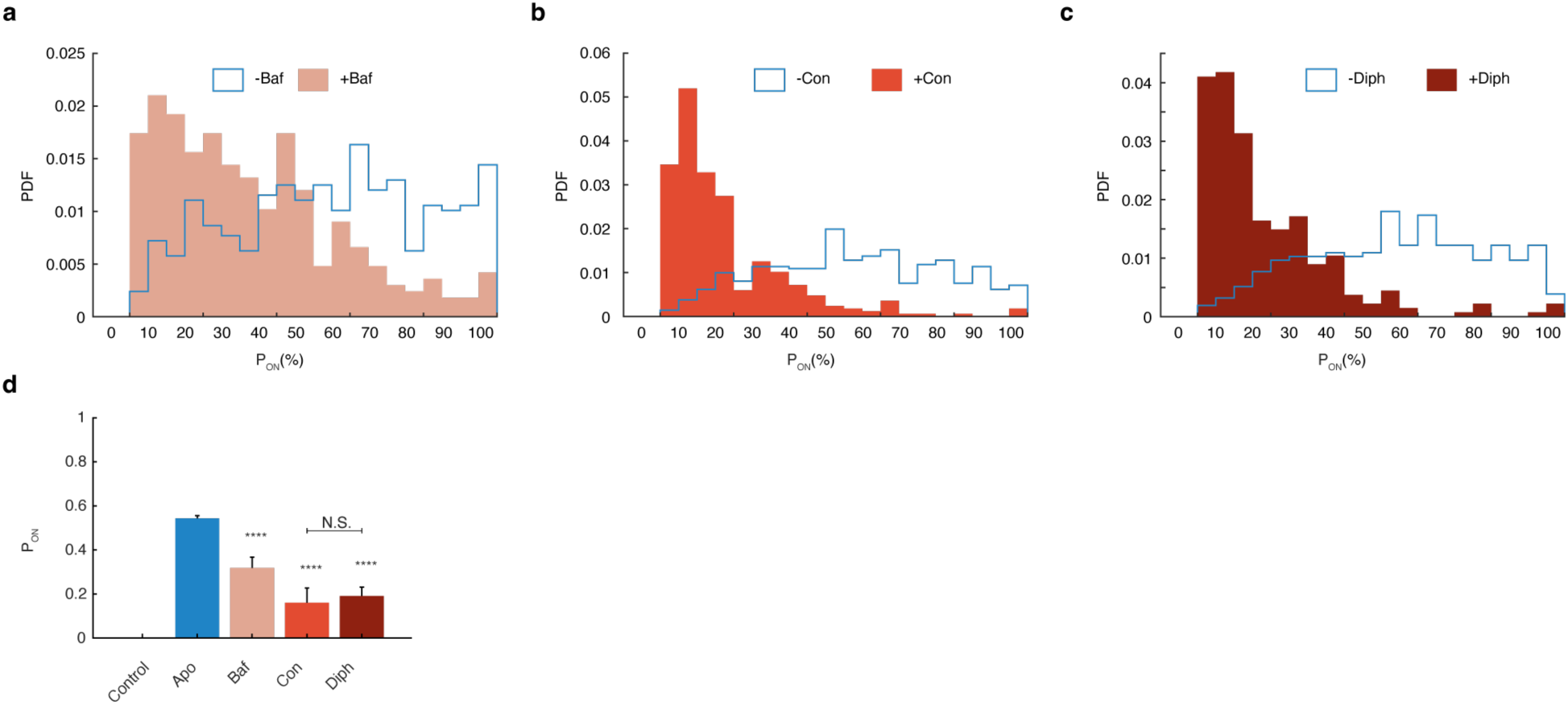
Kinetic experiments demonstrate that all inhibitors regulate the single-molecule active mode probability. **a-c**, Distributions of single-molecule active mode probability in the presence and absence of inhibitors. All inhibitors reduced the probability that the V-ATPase is found in an active mode. Concentrations of the inhibitors used were: 50 pM Baf, 320 pM Con, and 50 nM Diph. \*\*\*\**P* < 10^-10^ (± Baf); \*\*\*\**P* < 10^-10^ (± Con); **** *P* < 10^-10^ (± Diph); two-sided two-sample Kolmogorov-Smirnov test. **d**, Bar charts of the regulation of active mode probability (P_ON_). All three inhibitors investigated significantly reduced the probability of the V-ATPase being found in an active mode. mean ± s.d. from independent experiments; n.s. (*P* = 0.20, Con-Diph); two-sided two-sample Kolmogorov-Smirnov test; *n* = 11 (*apo*); *n = 3* (Baf); *n* = 4 (Con, Diph);

**Extended Data Fig. 8.**
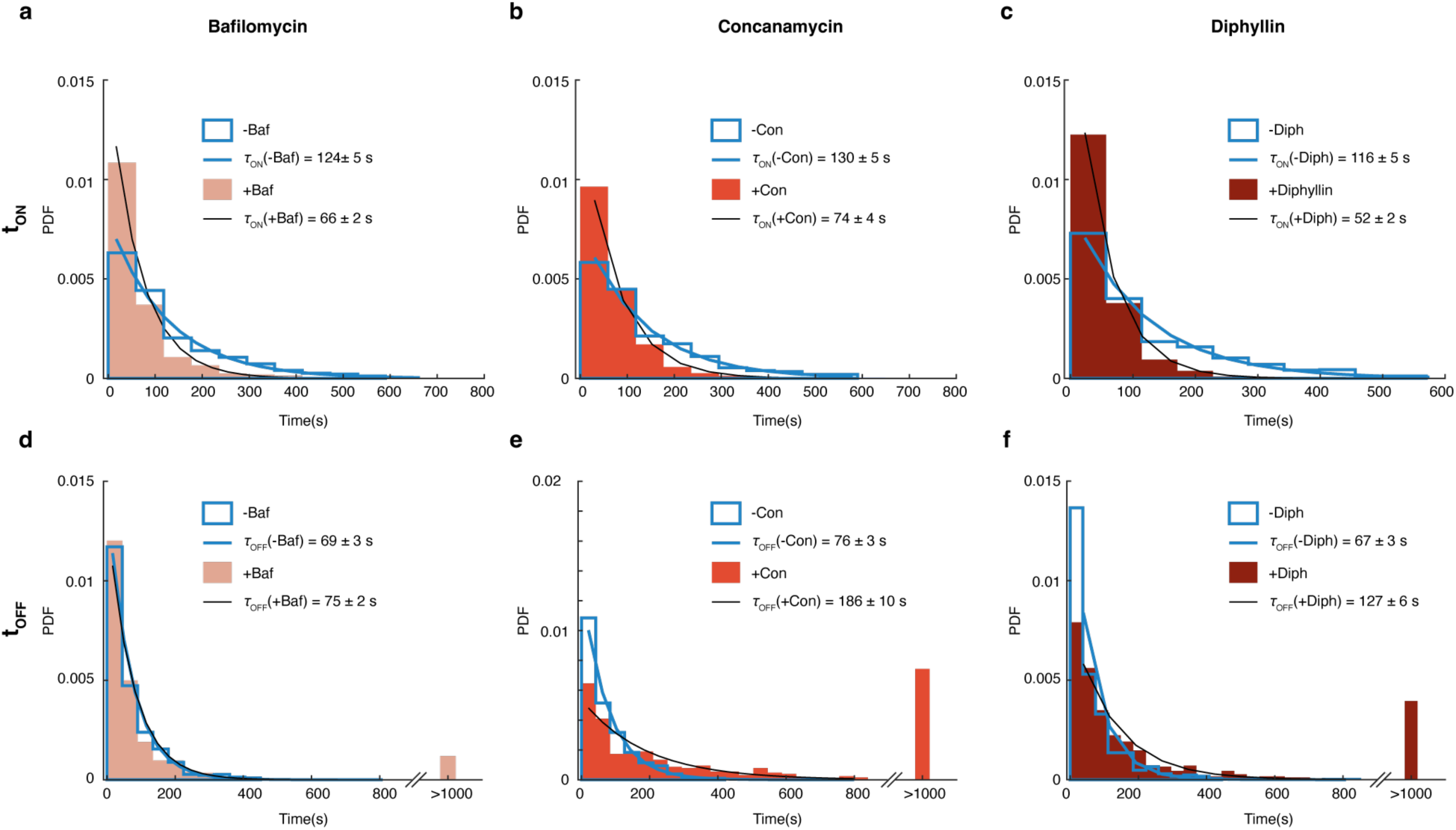
Bafilomycin regulates active mode dwell times but leaves the inactive mode unaffected; Concanamycin and Diphyllin both simultaneously downregulate active mode dwell times while upregulating inactive mode dwell times. **a-f**, Active and inactive mode dwell times in the presence and absence of inhibitors. Colored histograms correspond to dwell time populations in the presence of inhibitory compounds, while outlined histograms correspond to dwell times in the presence of only ATP. Characteristic dwell times (*τ*_ON_ and *τ*_OFF_) were extracted by fitting histograms with single exponentials. Errors correspond to the standard error of the fit.

**Extended Data Table 1.** Calculations of forward, backward transition activation barriers and free energy differences between active and inactive modes in the absence (*apo*) and presence of inhibitors. We employed transition state theory to derive the forward (ΔG^‡^_*ON*_) and backward (ΔG^‡^_*OFF*_) activation barriers, as well as the free energy difference (ΔG). For more details, we refer the reader to the corresponding section in the Supplementary discussion. The concentrations of the inhibitors were 50 pM (Baf), 320 pM (Con), and 50 nM (Diph).

| | $\Delta G^{\ddagger}_{ON}(\text{kJ} \cdot \text{mol}^{-1})$ | $\Delta G^{\ddagger}_{OFF}(\text{kJ} \cdot \text{mol}^{-1})$ | $\Delta G(\text{kJ} \cdot \text{mol}^{-1})$ |
| --- | --- | --- | --- |
| *Apo* | $84.9 \pm 0.1$ | $83.5 \pm 0.1$ | $-0.4 \pm 0.1$ |
| Bafilomycin | $83.4 \pm 0.1$ | $83.7 \pm 0.1$ | $1.9 \pm 0.6$ |
| Concanamycin | $83.6 \pm 0.1$ | $85.9 \pm 0.1$ | $4.1 \pm 1.2$ |
| Diphyllin | $82.7 \pm 0.1$ | $85.0 \pm 0.1$ | $3.6 \pm 0.7$ |

## Acknowledgements

We thank M. Grabe for providing the MatLab code for fitting the non-equilibrium model. We thank K. Huber for inspiring discussions that lead to the inception of this project. This work was supported by the Novo Nordisk Foundation (NNF17OC0028176) and the Lundbeck Foundation (Professorship grant R441-2023-360). R.J. was supported by an ERC Advanced Grant (SVNeuroTrans).

## Author contributions

D.S. was responsible for project management and for supervision. D.S. conceived the strategy with the help of E.K. E.K and D.S. designed research. E.K. developed the single-molecule assay and performed all experiments with the help of M.F who performed several controls. E.K. performed data analysis with the help of M.F. who analysed control experiments and M.C.I who developed the statistical analysis of dwell time distributions and corresponding simulations. C.G.S. was the principal software developer. J.P. prepared all the biochemical samples. P.J.J. and J.L.P. optimized the stochastic event detection algorithm. D.S. and E.K wrote the main text. E.K. prepared all figures and supplementary information. All authors discussed the results and commented on the manuscript.

## Competing interests

D.S. is the founder of Atomos Biotech. The other authors declare no competing interests.

## Supplementary discussion

### Intensity plateaus and step counting

We have shown previously that the presence of multiple transporters in a single vesicle would appear as multiple steps in the acidification plateaus^8^. Indeed, performing a step analysis we found that 85% ± 3% of the single vesicle data showed a single step, 13% ± 2% showed 2 steps and 2% ± 1% showed 3 steps or more (Supplementary Fig. 3). As a result, our data suggests that, on average, the SV contains 1.2 ± 0.1 copies of active V-ATPases which is in excellent agreement with proteomic studies^27–29^. These findings confirm the intrinsic single-molecule nature of the SV. All single-molecule analysis was restricted to single-step data.

### pH gradients at the thermodynamic limit

To estimate the maximum pH gradient that a V-ATPase can establish, we considered the free energy change of ATP hydrolysis under various intracellular conditions. The V-ATPase is assumed to translocate 10 protons per 3 ATP molecules hydrolyzed.

The Gibbs free energy change for ATP hydrolysis is given by:

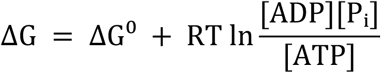

where ΔG⁰ = –30 kJ·mol⁻¹, R = 8.314 × 10⁻S kJ·mol⁻¹·K⁻¹, and T = 298 K. The free energy available per proton translocated is:

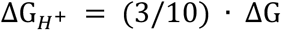

This energy drives the formation of a transmembrane pH gradient, where:

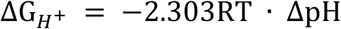

Solving for the pH gradient gives:

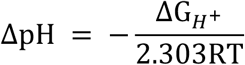

Using a fixed ATP concentration of 1 mM and 1 μM ADP and 1 μM Phosphate (in our experiments, the concentrations of P_i_ and ADP are likely much lower), we computed the theoretical maximum to be ΔpH_max_ = 4.28. These calculations define the thermodynamic upper limit of proton motive force generation by V-ATPases and demonstrate that in our single-molecule experiments, the gradient generated by the V-ATPase is substantially below the thermodynamic limit (ΔpH_SM_ ∼ 1)

### Kinetic assay: dwell time analysis

Employing the stochastic filtering algorithm, mode-switching events in single-molecule data were identified, and manual fine-tuning of events was performed on approximately 30-40% of the detected events.

Dwell time calculations considered only single-molecule data exhibiting at least one stochastic event. Active dwell times (t_ON_) were determined from the onset of acidification to the point at which the signal decreased, signifying V-ATPase deactivation. Inactive dwell times (t_OFF_) were computed from the start of signal decrease until the subsequent active event. Analysis of dwell times commenced from the initiation of the first proton-pumping event, uninterrupted by the recording’s conclusion.

### Kinetic assay: probability estimates

In kinetic experiments, the probability that the V-ATPase is found in an active mode is estimated as, *P*_ON_ = ∑ *t*_ON_ /*t*_*treatment*_, where *t*_*treatment*_ is the total duration of the recording under a specific condition (e.g., in the presence of an inhibitor).

### Snapshot assay: probability estimates and ensemble average acidification

The number of active vesicles for each treatment (*i*) is normalized to the total number of active vesicles identified under all conditions. We calculate the uncorrected probability of the active mode per treatment to be

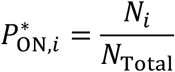

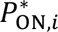 includes SVs that contain more than one V-ATPase and therefore skew the total probability towards higher active mode probability. To correct the probability for the number of active V-ATPases, we perform the following correction:

The probability that a SV with *n* V-ATPases, ^*n*^*P*_*OFF*_, is found in an inactive mode is given by

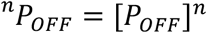

Where, *P*_*OFF*_is the inactive mode probability (*P*_*OFF*_ = 1 − *P*_*ON*_) of a SV containing a single V-ATPase. For the entire population of SVs then the total probability to be found in an inactive mode becomes

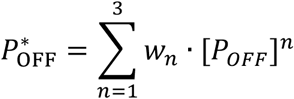

Where *w*_*n*_ is the corresponding fraction of SVs containing *n* = 1, 2 or 3 (or more) V-ATPases as calculated by the step analysis (Supplementary discussion and Supp. Fig. 4). Therefore, the corrected active mode probability, *P*_*ON*_, is numerically extracted using

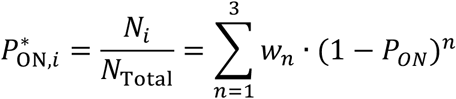

The average acidification per treatment *i,* (ΔpH_AV,i_) is a convolution of the single-molecule acidification of that treatment (ΔpH_SM,i_) reduced by the probability to be found in an active mode (*P*_ON_) and is expressed as

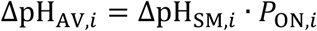

### Physical model

We employed the previously developed non-equilibrium model to extract pumping rates and permeabilities from single-molecule data^8,9,32^. The following system of equations describes the acidification kinetics of single synaptic vesicles by the V-ATPase:

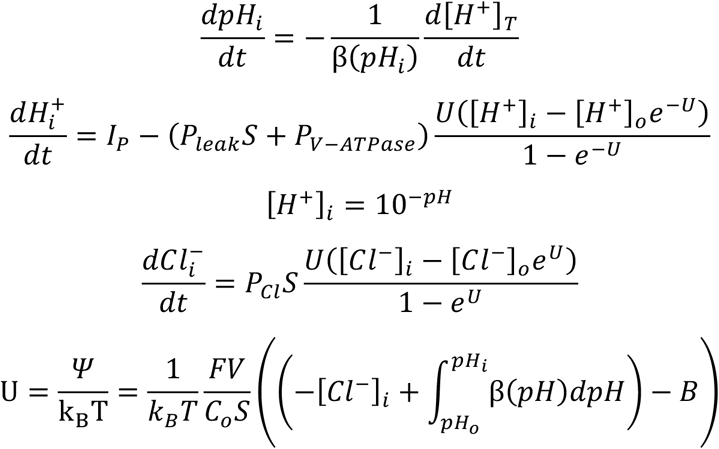

Concentrations of ionic species are represented as quantities in brackets. Subscripts *i*, *o* indicate intraluminal and extraluminal values of variables, respectively. Luminal buffering capacity is defined as *β*. In the lumen of SVs, protons are buffered by MOPS and pHrodo-DOPE molecules (the buffering capacity of the SV proteome has not been considered). Single-molecule proton pumping rates, passive proton permeability, and transprotein permeability are represented by *I_p_*, *P_leak_,* and *P_V-ATPase_* respectively and are the free parameters in the model. *S* and *V* are the surface area and volume of SVs, respectively. P_Cl_ is the permeability of SVs to *Cl^-^* and is chosen such that chloride ions can traverse freely across the membrane. *Ψ* is the membrane. Single-molecule traces fitted with the model are shown in E.D. Fig. X.

In a previous publication, we demonstrated the existence of a transprotein leakage mode of the V-ATPase. To fit the single-molecule data with the physical model, it was necessary to include transprotein permeability as a free parameter. However, in the present manuscript, the leaky mode was not further investigated.

### Calculation of the transition energy barrier heights for on-to-off and off-to-on cycle transitions

Transition rates (main Fig. 3c, E.D. Fig. 6) were calculated using transition state theory^44^, where the pumping and inactive modes are considered ground states separated by an activation barrier (the transition state). According to transition state theory, the energy required to achieve the transition state (and cross the barrier) is:

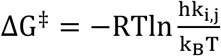

where R is the gas constant, T is the absolute temperature, k_B_ is Boltzmann’s constant, h is Planck’s constant, k_i,j_ is the measured transition rate from active to inactive or from inactive to active modes. Under the assumption that we have a two-mode model, active and inactive mode rate constants can be calculated using equations

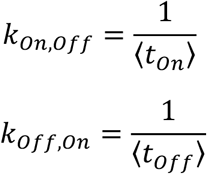

Where, 〈*t*_*on*_〉 and 〈*t*_*off*_〉 are the average dwell times in active and inactive modes for the V-ATPase.

### Calculation of free energy difference between active and inactive modes

We calculated the Gibbs free energy difference between active and inactive modes of the V-ATPase for the *apo* and in the presence of inhibitors (see Fig. 6 and E.D. Table 1) using the equation,

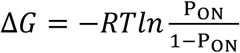

where P_ON_ and 1-P_ON_ represent the probabilities of the pump being in active or inactive modes, respectively.

### Stochastic model description

Here, we provide a brief summary of the Bayesian event detection methodology developed in a previous work. For a more detailed description of the method, we refer the reader to an earlier publication^8^.

The signal in the experiment refers to variations in light intensity caused by the activity of V-ATPase. This signal depends on change-point times ***T*** = {*T*_1_, *T*_2_, …, *T*_*n*_} representing mode-switching between active and inactive modes of the proton pump. The proton-pumping mode leads to increased fluorescence modelled as an exponential increase to an equilibrium level (α) with rate (β), while the inactive mode results in exponential decay back to baseline with rate (γ). The overall signal model, μ(t), switches between these modes at change-points.

Exponential rates for proton-pumping and inactive events are denoted as β*_i_* and γ*_i_*, respectively. Equilibrium level α remains constant. Parameters are grouped as 𝝂 = {*α*, *β*_1_, *γ*_1_, …, *β*_*n*_, *γ*_*n*_}. Time between mode-switching events is denoted as ***θ*** = {*θ*_1_, *θ*_2_, …, *θ*_*n*_} = {*T*_1_, *T*_2_ − *T*_1_, …, *T*_*n*_ − *T*_*n*–1_}, which replaces ***T*** in the model. Unknowns, including change-point times, are treated as random variables with unbiased prior distributions in a Bayesian setting.

Stochastic filtering, based on Bayesian techniques, is employed to estimate the signal, considering prior opinions and updating with observed data. Maximum a posteriori probability (MAP) estimates for change-points and parameters are obtained from the joint distribution. The likelihood function *L* is defined based on discrete observed data, approximating continuous-time likelihood. Numerical constraints are applied for parameter ranges.

A sliding method of Bayesian stochastic filtering is introduced to address the computational complexity associated with a large number of change-points. This method segments data into regions with one mode-switch event, reducing complexity. The Bayesian event detection algorithm is novel, focusing on raw data to identify mode-switching events without smoothing.

### Model selection via simulated Bayesian Information Criterion (BIC) differences

To determine whether lifetime distributions are better described by a single or double-exponential model, we performed fitting and simulation-based inference. First, lifetimes were fit using both a single- and double-exponential model for maximum likelihood estimation (MLE) with grid search for ensuring global maxima to obtain parameter estimates. The Bayesian Information Criterion (BIC) is defined as

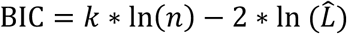

and the change in BIC as

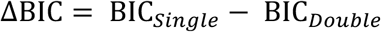

where indices indicate BIC values for a single- and double-exponential model respectively. *k* describes the number of fit parameters for the respective model, *n* the number of events and *L̂* the maximized likelihood value, which in our case is extracted from the best fit.

Next, we generated 500 synthetic datasets under each model given the fit results and data size of the experiment. We then performed the same MLE on each synthetic dataset with both a single- and double-exponential model and determined their ΔBIC.

We then estimated the probability density functions (pdf) of observing a given ΔBIC value under each model, p_Single_(ΔBIC) and p_Double_(ΔBIC) respectively using Kernel Density Estimation (KDE) in Python^45^.

The posterior probability of a double-exponential distribution producing the experimental ΔBIC value (termed ΔBIC*) was then calculated using Bayes theorem via

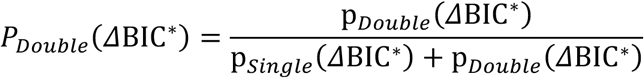

assuming equal priors since no prior knowledge is available to favor either of the two models.

### Supplementary Methods

#### Image analysis

Data from light microscopy experiments were analyzed using a custom software suite and MATLAB scripts. A single particle localization algorithm^46^ identified vesicles and their positions, with a drift correction algorithm^47^ addressing potential XY drifts caused by thermal fluctuations. Following image stabilization and particle identification, 3x3 pixel regions of interest (ROIs) were defined around each particle, and integrated intensity readouts were obtained by summing the pixel values.

Single-molecule normalized intensity, defined as raw intensity divided by the mean baseline raw intensity (*I*_SM_ *= I*_raw_*/I*_base_), and standardized intensity, defined as raw intensity reduced by the mean baseline intensity and divided by the standard deviation of the signal at the baseline (Standardized Intensity *= (I*_raw_*-I*_base_*)/σ*_base_), were employed. Normalized intensities were used for data analysis and visualization, as well as to convert single-molecule traces to pH units and classify traces based on their number of intensity steps (E.D. Fig. X and Y). Standardized intensities were used in the stochastic filtering algorithm.

### Snapshot assay for compound screening

To screen compounds using the snapshot assay, we first acquire a stack of images of immobilized vesicles in activity buffer (300 mM glycine, 2 mM MOPS, 2 mM MgSO₄, pH 7.1–7.15, adjusted with Tris). This defines the baseline fluorescence of synaptic vesicles (SVs).

Next, 1 mM MgATP is added to initiate proton pumping, along with 30 mM choline chloride to counteract positive charge accumulation in the SV lumen^22,48,49^. This condition reveals vesicles that respond to ATP, indicating constitutively active V-ATPase; this is referred to as the *apo* condition.

Subsequently, increasing concentrations of V-ATPase inhibitors are sequentially introduced at 6.5-minute intervals, allowing incubation. After each interval, a 20-frame stack is acquired.

Active vesicles were identified for each treatment by thresholding based on the intensity change relative to the baseline. A trace was classified as active if ≥50% of frames exceeded the threshold. For each treatment, the number of active traces and mean fluorescence intensity per vesicle were extracted from the recording. Mean intensities were converted to ΔpH values and used to estimate acidification and proton pumping rates.

At the end of the imaging sequence, a 20-frame stack was acquired under saturating pH conditions to calibrate fluorescence-to-pH conversion (Supplementary Fig. 2, Supplementary Methods).

For contour plots and population histograms (Fig. 2–4, Extended Data Fig. 3), intensities were normalized per vesicle, such that the baseline/inactive mode was set to 0 and the active mode was set to 1. This normalization preserves intra-vesicle variability across conditions. Mode-occupancy distributions were fitted with double Gaussians via maximum likelihood estimation with grid search for ensuring global maxima. When active-mode occupancy was too low for robust fitting, a lower bound was imposed on the mean of that population.

Note: All snapshot experiments used 1 mM ATP supplemented with 30 mM choline chloride unless stated otherwise.

### Single-vesicle kinetic assay

Acidification kinetics were initiated by recording baseline fluorescence of immobilized vesicles in activity buffer (300 mM glycine, 2 mM MOPS, 2 mM MgSO₄, pH 7.1–7.15, adjusted with Tris) for 5 minutes. Activity was then triggered by adding 1 mM MgATP and 30 mM choline chloride to counteract the buildup of membrane potential^22,48,49^. After a 10-minute equilibration, compounds were introduced, and vesicle responses were monitored over the subsequent 15 minutes. At the end of the recording, a 20-frame image stack was acquired under saturating pH conditions to enable fluorescence-to-pH conversion via calibration curves (Supplementary Fig. 2, Supplementary Methods).

Active V-ATPase transport kinetics were identified using signal-to-noise ratio (SNR)– based thresholding. SNR for each vesicle was defined as

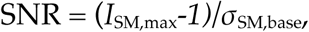

where *I*_SM,max_ is the peak normalized intensity, and *σ_SM,base_* is the baseline standard deviation. To reduce spike-related outliers in *I*_SM,max_, traces were smoothed with a Chung– Kennedy filter^50^.

Active transport traces were manually classified based on the number of discrete intensity steps (plateaus; Supplementary Fig. 3). Only vesicles exhibiting a single step— indicative of individual V-ATPase activity—were included in the single-molecule analysis. Traces lacking characteristic stochastic behaviour were excluded.

Note: All kinetic experiments included 1 mM ATP and 30 mM choline chloride unless otherwise specified.

### Calibration of pH in vesicles

A local calibration approach was employed using DOPE-pHrodo labeled liposomes with the specified lipid composition (DOPC:DOPS:Chol:(18:1 Biotinyl Cap PE):DOPE-pHrodo 64.4:10:25:0.5:0.1). Immobilized liposomes were subjected to calibration buffer (150 mM K-Gluconate - 2 mM MOPS - 2 mM MgSO_4_) with 60 nM valinomycin and 5 μM CCCP at nine pH values. The custom software identified vesicles, and raw intensities were extracted and fitted with a sigmoidal function. Normalization to pH 7.1 baseline intensity yielded mean values for pKa and rates. *I*_base_ and *I*_max_, the defining fluorophore dynamic range, were locally defined for each vesicle after activity acquisition.

Following single vesicle acidification kinetics at pH 7.10-7.15, we incubated SVs in saturation buffer (300 mM Glycine - 2 mM MOPS – 2 mM MgSO4) at pH 2.85-3.15 (adjusted with citric acid) with 5 μM CCCP and 30 mM choline chloride. Calibration involved normalizing raw traces, accounting for pHrodo-DOPE bilayer contribution, and calculating pH as a function of intensity. The assumption of equal distribution of pHrodo-DOPE lipids between bilayer leaflets was maintained throughout the analysis. For a more detailed description of the calibration procedure, we refer the reader to previous work^8^.

### Single SV pH gradients

To extract single SV acidification equilibria (ΔpH_SM,max_) we selected regions of the kinetic traces during which proton pumping and proton leakage are in equilibrium. In these regions, the pH gradient established by the V-ATPase is the maximal possible given that the permeability of the membrane counteracts the proton pumping. For a region of a trace to qualify to extract the ΔpH_SM,max_, it must display a stable plateau for at least 5 frames (typically 15 seconds of real-time imaging). The value of ΔpH_SM,max_ is the mean value of the frames in the plateau selected.

---

**Supplementary Fig. 1.**
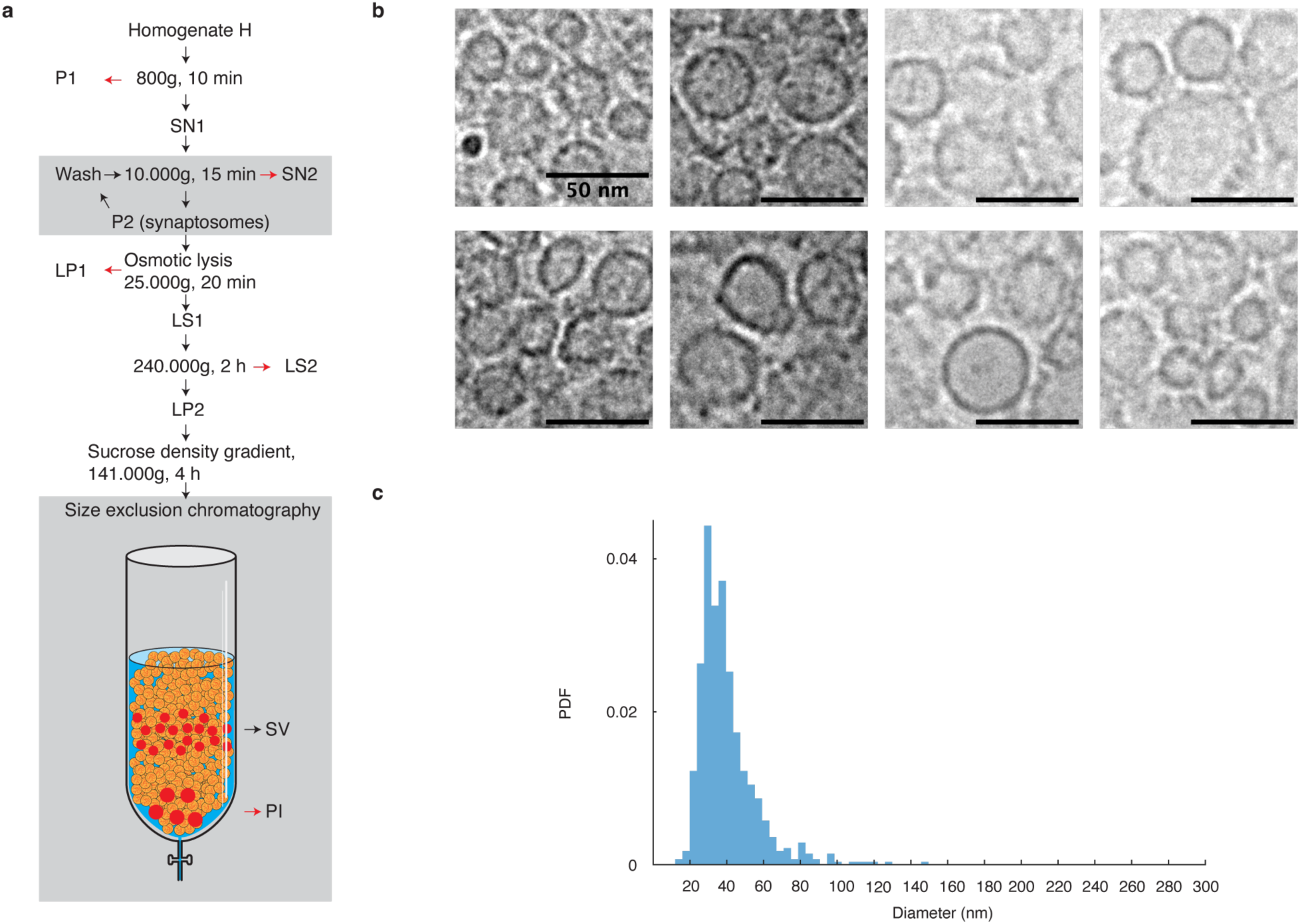
Purification and morphology of SVs. **a**, Schematic illustration of the purification protocol for synaptic vesicles (SVs). **b**, Cryo-electron micrographs depicting the high purity of the SV fraction. Larger vesicles were infrequently observed in the images. The data is derived from numerous micrographs obtained during a single experimental session. The scale bar corresponds to 50 nm. **c**, Size distribution of SVs as observed in cryo-electron micrographs. The SV population displayed homogeneous morphology and a uniform distribution, with a consistent mean vesicle diameter of 38 ± 1 nm based on the S.E. of the fit.

**Supplementary Fig. 2.**
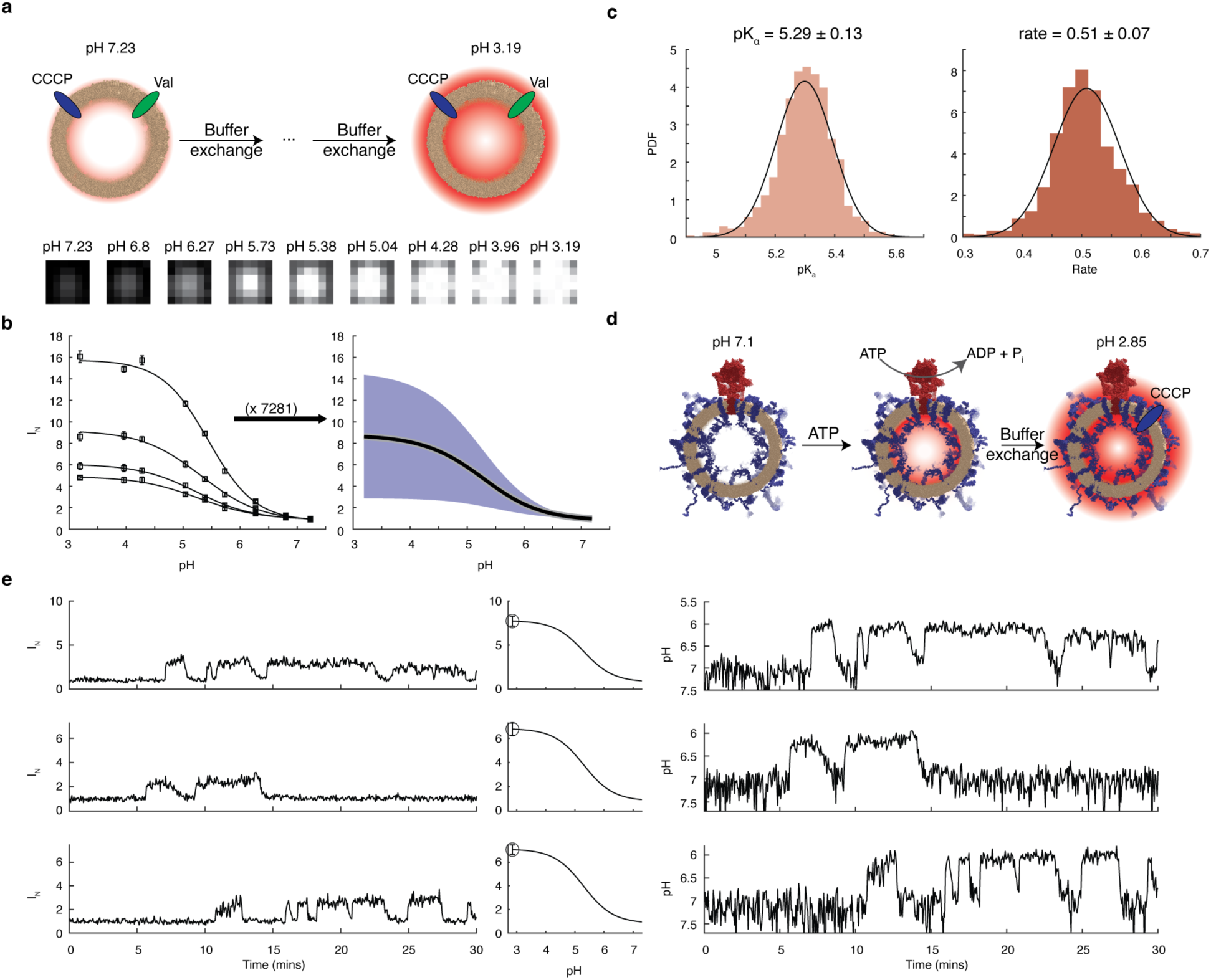
Single-vesicle-specific pH calibration. **a**, Illustration of the pH titration experiments to determine the calibration parameters pK_a_ and growth rate. Briefly, large unilamellar vesicles (LUVs) are immobilized on the glass surface and incubated in a calibration buffer containing K-Gluconate-MOPS at pH 7.23. Then buffers of the same composition with decreasing pH are sequentially exchanged in the presence of CCCP and valinomycin to maintain ionic balance and negate the membrane potential across vesicular membranes. **b**, Left: example data from single-vesicle pH titrations that are fitted with a sigmoidal function. Data points and error bars correspond to the mean and s.d. of stacks of 20 images per data point. Right: ensemble average (mean) and spread (s.d.) of the sigmoidal fits to the total population of LUVs. **c**, Distributions of the dissociation constant, pK_a_, and growth rate from single LUV fits. **d**, Schematic of a typical single-molecule activity measurement. SVs are immobilized on glass, and subsequently, ATP is injected to commence activity. At the end of the measurement, a saturation buffer containing glycine-MOPS at a pH of ∼3 is injected, along with CCCP and choline chloride, to neutralize the membrane potential. A stack of 20 images is then captured. This stack is used to calibrate intensities to pH as described briefly in the supplementary discussion and Ref. ^8^. **e**, Example traces before (left) and after (right) calibration along with their corresponding calibration curve (mid). Each calibration curve is created using the dissociation constant and the growth rate determined in c, as well as the saturation and baseline intensity determined in situ from activity and saturation measurements, which are unique for each LUV.

**Supplementary Fig. 3.**
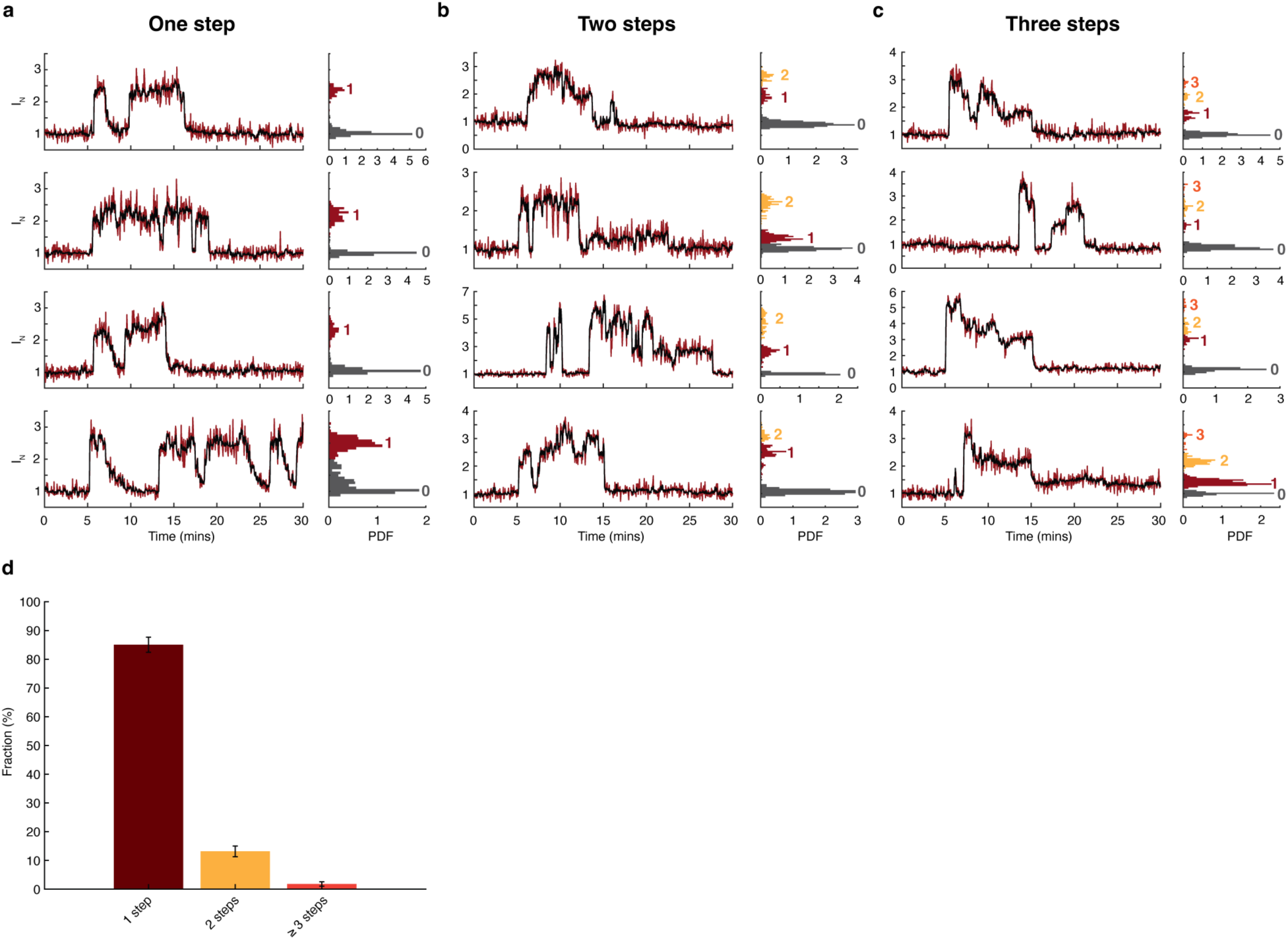
Synaptic vesicles contain predominantly single V-ATPases. **a-c**, Left: Single vesicle acidification kinetics showcasing mode-switching dynamics with different acidification plateaus (steps). Each acidification plateau corresponds to a different dynamic equilibrium between pumping and leakage. Red traces correspond to normalized intensities, while black traces correspond to data treated with a Chung-Kennedy filter. Right: Histograms of accumulated intensities for single-vesicle traces. Distinct populations of steps can be identified. **a**, Representative traces showcasing a single acidification plateau. A single plateau corresponds to a single V-ATPase switching to an active mode. **b**, Representative traces that contain two plateaus correspond to two V-ATPases stochastically switching on. **c**, Similarly, traces that contain three or more plateaus correspond to multiple V-ATPases switching to an active mode in a single SV. **d**, Bar chart of the fraction of vesicles that showed single (dark red), double (yellow), or multiple plateaus (orange). Taken together, on average, the synaptic vesicle contains 1.2 ± 0.1 V-ATPases; *n* = 3.

**Supplementary Fig. 4.**
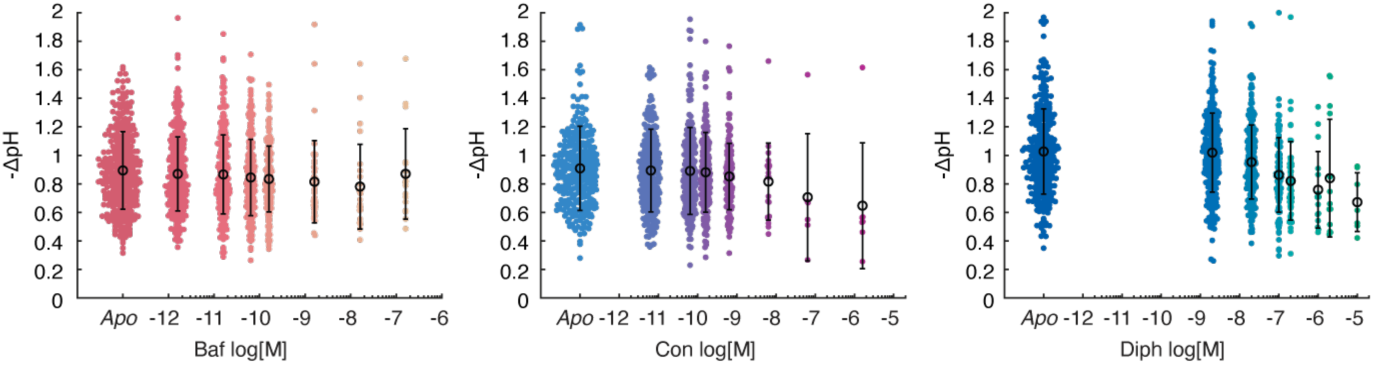
Inhibitor titrations do not regulate acidification levels but the fraction of V-ATPases in an active mode. Beeswarm plots of single-molecule acidification as a function of inhibitor concentration for 50 pM bafilomycin, 320 pM concanamycin, and 50 nM diphyllin. Coloured data points correspond to ΔpH_SM_ by individual SVs, while black data points and error bars correspond to the mean and one s.d. of the population of SVs. ΔpH_SM_,. Remarkably, the number of active SVs is markedly reduced in the presence of increasing concentrations of inhibitors. Taken together, all inhibitors predominantly regulate the probability that the V-ATPase enters inactive modes rather than regulating ΔpH_SM_. Data has not been corrected for the control.

**Supplementary Table 1.** Bayesian analysis reveals that bafilomycin introduces a second exponential lifetime for the active mode while concanamycin and diphyllin modulate the single exponential lifetime of the *apo* condition. All inhibitors stabilized a secondary inactive mode. As described in supplementary discussion, modelling of the difference of the Bayesian Information Criterion (ΔBIC) distributions for single and double exponential fits of lifetime distributions determined the probability that the lifetimes are expressed by a double exponential for the active, 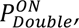 and the inactive, 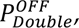 mode. Active mode: the *apo* condition is best represented by a single exponential distribution as the double exponential Maximum Likelihood Estimation (MLE) based fit generated two monoexponentials with identical lifetimes. Bafilomycin, unambiguously introduced a second population of ON lifetimes. Concanamycin and diphyllin data are best represented by a single exponential. Inactive mode: we cannot confirm whether the *apo* condition is best represented by a single or a double exponential as 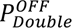 is not significant. However, all inhibitors unambiguously stabilized a secondary exponential with high statistical significance. ^‡^We fitted ON lifetimes of concanamycin and diphyllin with a double exponential model that generated exponential terms with overwhelming relative weight of one term over the other (98% for Con and >99% for Diph). Therefore, both datasets are defined as single exponentials.

|  | *Apo* | Baf | ConA | Diph |
| --- | --- | --- | --- | --- |
| $P_{Double}^{ON}$ | 0 (n.s.) | >0.99 (**) | 0<sup>‡</sup> (n.s.) | 0<sup>‡</sup> (n.s.) |
| $P_{Double}^{OFF}$ | 0.69 (n.s.) | >0.99 (**) | 0.99 (**) | 0.98 (*) |

## Notes

### Competing Interest Statement

Dimitrios Stamou is the founder of Atomos Biotech. The other authors declare no competing interests.

## References

1. Nishi, T. & Forgac, M. The vacuolar (H+)-ATPases — nature’s most versatile proton pumps. Nat. Rev. Mol. Cell Biol. 3, 94–103 (2002).

2. Vasanthakumar, T. & Rubinstein, J. L. Structure and Roles of V-type ATPases. Trends Biochem. Sci. 45, 295–307 (2020).

3. Chen, F., Kang, R., Liu, J. & Tang, D. The V-ATPases in cancer and cell death. Cancer Gene Ther. 29, 1529–1541 (2022).

4. Collins, M. P. & Forgac, M. Regulation and function of V-ATPases in physiology and disease. Biochim. Biophys. Acta BBA - Biomembr. 1862, 183341 (2020).

5. Wang, R. et al. Molecular basis of V-ATPase inhibition by bafilomycin A1. Nat. Commun. 12, 1782 (2021).

6. Keon, K. A., Benlekbir, S., Kirsch, S. H., Müller, R. & Rubinstein, J. L. Cryo-EM of the Yeast V_O_ Complex Reveals Distinct Binding Sites for Macrolide V-ATPase Inhibitors. ACS Chem. Biol. 17, 619–628 (2022).

7. Bowman, B. J., McCall, M. E., Baertsch, R. & Bowman, E. J. A Model for the Proteolipid Ring and Bafilomycin/Concanamycin-binding Site in the Vacuolar ATPase of Neurospora crassa. J. Biol. Chem. 281, 31885–31893 (2006).

8. Kosmidis, E. et al. Regulation of the mammalian-brain V-ATPase through ultraslow mode-switching. Nature 611, 827–834 (2022).

9. Veshaguri, S. et al. Direct observation of proton pumping by a eukaryotic P-type ATPase. Science 351, 1469–1473 (2016).

10. Akyuz, N., Altman, R. B., Blanchard, S. C. & Boudker, O. Transport dynamics in a glutamate transporter homologue. Nature 502, 114–118 (2013).

11. Akyuz, N. et al. Transport domain unlocking sets the uptake rate of an aspartate transporter. Nature 518, 68–73 (2015).

12. Fitzgerald, G. A. et al. Quantifying secondary transport at single-molecule resolution. Nature 575, 528–534 (2019).

13. Ciftci, D. et al. Single-molecule transport kinetics of a glutamate transporter homolog shows static disorder. Sci. Adv. 6, eaaz1949 (2020).

14. Furuike, S. et al. Resolving stepping rotation in Thermus thermophilus H+-ATPase/synthase with an essentially drag-free probe. Nat. Commun. 2, 233 (2011).

15. Arai, S. et al. Rotation mechanism of Enterococcus hirae V1-ATPase based on asymmetric crystal structures. Nature 493, 703–707 (2013).

16. Abbas, Y. M., Wu, D., Bueler, S. A., Robinson, C. V. & Rubinstein, J. L. Structure of V-ATPase from the mammalian brain. Science 367, 1240–1246 (2020).

17. Yuan, F.-L. et al. The vacuolar ATPase in bone cells: a potential therapeutic target in osteoporosis. Mol. Biol. Rep. 37, 3561–3566 (2010).

18. Dowdy, S. F. Overcoming cellular barriers for RNA therapeutics. Nat. Biotechnol. 35, 222–229 (2017).

19. Petros, R. A. & DeSimone, J. M. Strategies in the design of nanoparticles for therapeutic applications. Nat. Rev. Drug Discov. 9, 615–627 (2010).

20. Eaton, A. F., Merkulova, M. & Brown, D. The H^+^ -ATPase (V-ATPase): from proton pump to signaling complex in health and disease. Am. J. Physiol.-Cell Physiol. 320, C392–C414 (2021).

21. Ahmed, S., Holt, M., Riedel, D. & Jahn, R. Small-scale isolation of synaptic vesicles from mammalian brain. Nat. Protoc. 8, 998–1009 (2013).

22. Farsi, Z. et al. Single-vesicle imaging reveals different transport mechanisms between glutamatergic and GABAergic vesicles. Science 351, 981–984 (2016).

23. Stamou, D., Duschl, C., Delamarche, E. & Vogel, H. Self-Assembled Microarrays of Attoliter Molecular Vessels. Angew. Chem. Int. Ed. 42, 5580–5583 (2003).

24. Bendix, P. M., Pedersen, M. S. & Stamou, D. Quantification of nano-scale intermembrane contact areas by using fluorescence resonance energy transfer. Proc. Natl. Acad. Sci. 106, 12341–12346 (2009).

25. Mathiasen, S. et al. Nanoscale high-content analysis using compositional heterogeneities of single proteoliposomes. Nat. Methods 11, 931–934 (2014).

26. Tonnesen, A., Christensen, S. M., Tkach, V. & Stamou, D. Geometrical Membrane Curvature as an Allosteric Regulator of Membrane Protein Structure and Function. Biophys. J. 106, 201–209 (2014).

27. Wang, C. et al. Structure and topography of the synaptic V-ATPase–synaptophysin complex. Nature 1–6 (2024) doi:10.1038/s41586-024-07610-x.

28. Takamori, S. et al. Molecular Anatomy of a Trafficking Organelle. Cell 127, 831–846 (2006).

29. Mutch, S. A. et al. Protein Quantification at the Single Vesicle Level Reveals That a Subset of Synaptic Vesicle Proteins Are Trafficked with High Precision. J. Neurosci. 31, 1461–1470 (2011).

30. Coupland, C. E. et al. High-resolution electron cryomicroscopy of V-ATPase in native synaptic vesicles. Science 385, 168–174 (2024).

31. Reddy, K. D., Ciftci, D., Scopelliti, A. J. & Boudker, O. The archaeal glutamate transporter homologue GltPh shows heterogeneous substrate binding. J. Gen. Physiol. 154, e202213131 (2022).

32. Ishida, Y., Nayak, S., Mindell, J. A. & Grabe, M. A model of lysosomal pH regulation. J. Gen. Physiol. 141, 705–720 (2013).

33. Bowman, E. J., Graham, L. A., Stevens, T. H. & Bowman, B. J. The Bafilomycin/Concanamycin Binding Site in Subunit c of the V-ATPases from Neurospora crassa and Saccharomyces cerevisiae. J. Biol. Chem. 279, 33131–33138 (2004).

34. Sørensen, M. G., Henriksen, K., Neutzsky-Wulff, A. V., Dziegiel, M. H. & Karsdal, M. A. Diphyllin, a Novel and Naturally Potent V-ATPase Inhibitor, Abrogates Acidification of the Osteoclastic Resorption Lacunae and Bone Resorption1*. J. Bone Miner. Res. 22, 1640–1648 (2007).

35. Hou, W. et al. Bioactivities and Mechanisms of Action of Diphyllin and Its Derivatives: A Comprehensive Systematic Review. Molecules 28, 7874 (2023).

36. Zosel, F., Mercadante, D., Nettels, D. & Schuler, B. A proline switch explains kinetic heterogeneity in a coupled folding and binding reaction. Nat. Commun. 9, 3332 (2018).

37. Greives, N. & Zhou, H.-X. Both protein dynamics and ligand concentration can shift the binding mechanism between conformational selection and induced fit. Proc. Natl. Acad. Sci. 111, 10197–10202 (2014).

38. Di Castro, M. A. et al. Local Ca2+ detection and modulation of synaptic release by astrocytes. Nat. Neurosci. 14, 1276–1284 (2011).

39. Zhang, Q., Li, Y. & Tsien, R. W. The Dynamic Control of Kiss-And-Run and Vesicular Reuse Probed with Single Nanoparticles. Science 323, 1448–1453 (2009).

40. Ziskin, J. L., Nishiyama, A., Rubio, M., Fukaya, M. & Bergles, D. E. Vesicular release of glutamate from unmyelinated axons in white matter. Nat. Neurosci. 10, 321–330 (2007).

## References

41. Kemmer, G. C. et al. Lipid-conjugated fluorescent pH sensors for monitoring pH changes in reconstituted membrane systems. The Analyst 140, 6313–6320 (2015).

42. Taoufiq, Z. et al. Hidden proteome of synaptic vesicles in the mammalian brain. Proc. Natl. Acad. Sci. 117, 33586–33596 (2020).

43. Upmanyu, N. et al. Colocalization of different neurotransmitter transporters on synaptic vesicles is sparse except for VGLUT1 and ZnT3. Neuron 110, 1483–1497.e7 (2022).

44. Laidler, K. J. & King, M. C. Development of transition-state theory. J. Phys. Chem. 87, 2657–2664 (1983).

45. Virtanen, P. et al. SciPy 1.0: fundamental algorithms for scientific computing in Python. Nat. Methods 17, 261–272 (2020).

46. Furst, E. M. Particle tracking with Matlab. (2015).

47. Guizar, M. Efficient subpixel image registration by cross-correlation (https://www.mathworks.com/matlabcentral/fileexchange/18401-efficient-subpixel-image-registration-by-cross-correlation), MATLAB Central File Exchange. Retrieved September 28, 2021. (2021).

48. Preobraschenski, J., Zander, J.-F., Suzuki, T., Ahnert-Hilger, G. & Jahn, R. Vesicular Glutamate Transporters Use Flexible Anion and Cation Binding Sites for Efficient Accumulation of Neurotransmitter. Neuron 84, 1287–1301 (2014).

49. Preobraschenski, J. et al. Dual and Direction-Selective Mechanisms of Phosphate Transport by the Vesicular Glutamate Transporter. Cell Rep. 23, 535–545 (2018).

50. Chung, S. H. & Kennedy, R. A. Forward-backward non-linear filtering technique for extracting small biological signals from noise. J. Neurosci. Methods 40, 71–86 (1991).

